# Seasonal dynamics and sun/shade heterogeneity of leaf gas exchange and VOC emissions inside a tall temperate forest canopy

**DOI:** 10.64898/2026.01.23.701264

**Authors:** Stefanie Dumberger, Yasmina Frey, Clara Stock, Sophie Wehlings-Schmitz, Delon Wagner, Kathrin Kühnhammer, Lea Dedden, Markus Weiler, Markus Sulzer, Andreas Christen, Jürgen Kreuzwieser, Ulrike Wallrabe, Christiane Werner, Simon Haberstroh

## Abstract

Leaf gas exchange is the key driver of forest carbon uptake and directly determines forest carbon sink activity. Additionally, plants release a variety of biogenic volatile organic compounds (VOCs) acting as stress signals of trees. However, continuous hourly resolved measurements of leaf gas exchange and VOC emissions in tall tree canopies are challenging and remain scarce. To this end, we developed a sophisticated in-situ leaf gas exchange measurement system with 24 cuvettes deployed on mature *Fagus sylvatica* (n=3) and *Pseudotsuga menziesii* (n=3) individuals in a mixed temperate forest. We additionally measured sap flux density (*J_s_*), radial growth and tree water deficit (*TWD*) to gain a holistic picture of seasonal leaf and stem water and carbon flux dynamics during the summer of 2024.

During midsummer, we found a gradual reduction of stomatal conductance (*g_s_*) and VOC emissions of sun, but not shade branchlets of *P. menziesii* in response to moderate atmospheric and edaphic drying. Decreased *g_s_* led to a downregulation of transpiration (*E*), *J_s_*, and carbon isotope discrimination accompanied by an increase in *TWD* and intrinsic water used efficiency. Leaf gas exchange of shade branchlets remained unaffected due to microclimatic buffering effects. Contrarily, sun leaves of *F. sylvatica*, profited from sunny midsummer conditions and increased leaf gas exchange, whereas shade leaves benefitted from more diffuse light during early summer exhibiting similar carbon assimilation, transpiration and VOC emissions as sun leaves. For both species we found a clear time lag of four to five hours between maximum leaf and stem water fluxes and a delay of up to 20 hours for the recovery of *TWD*, highlighting the role of stem water reserves.

Pronounced seasonal and diurnal differences of leaf gas exchange, stem water fluxes and VOC emissions showed, that continuous data are essential to better understand variability of ecosystem flux dynamics.

## Introduction

Forests represent a major terrestrial carbon sink, taking up approximately 20% of annual anthropogenic CO_2_ emissions (Pan et al. 2024; Friedlingstein et al. 2025). Yet, increasing intensity of heat waves and droughts strongly compromise the carbon sink strength (Ciais et al. 2005; Schuldt et al. 2020; Gharun et al. 2024; Haberstroh et al. 2025; Knutzen et al. 2025; Werner et al. 2025). Leaf gas exchange is the key driver of carbon uptake and water loss in forest ecosystems shaping plant productivity and stress responses. Stomata jointly regulate carbon uptake and water release on the leaf level, which are both affected by microclimatic conditions (e.g. air temperature, solar irradiance, water availability, atmospheric humidity), and concentration gradients of CO_2_ and H_2_O between the atmosphere, the leaf boundary layer and the chloroplast (Martínez-Vilalta et al. 2014; Salmon et al. 2020). In response to drought stress, stomata close to prevent excessive water loss which has a cascading effect on intrinsic water use efficiency (*WUEi*) and leaf ^13^C discrimination (*Δ_leaf_)*. An increase in *WUEi* and a decrease in *Δ_leaf_* is expected in response to stomatal closure, yet the magnitude of change depends on the species-specific stress tolerance (Cernusak et al. 2013; Werner et al. 2021). Moreover, leaf exchange rates of biogenic volatile organic compounds (VOCs) via the stomata and their composition provide insights into stress status, defense mechanisms and communication strategies of trees (Laothawornkitkul et al. 2009). VOC emissions and compositions are species-specific and *de novo* synthesis is mainly controlled by microclimate and availability of fresh assimilates, yet several species including conifers also possess specialized storage compartments emitting VOCs dependent on air temperature and partially decoupled from synthesis (Lerdau et al. 1997; Niinemets et al. 2004a; Dindorf et al. 2006; Holzke et al. 2006; Joó et al. 2011; Van Meeningen et al. 2016). Under heat stress, VOC emissions are generally increased and some compounds, such as isoprene and monoterpenoids, are known to increase membrane stability and thermotolerance (Peñuelas and Llusià 2003; Copolovici et al. 2005; Werner et al. 2020; Bourtsoukidis et al. 2024; Meischner et al. 2024). Moderate drought stress induces an initial increase of VOC emissions, while under severe drought decreased emission rates in accordance with reduced photosynthetic rates were observed (Bertin et al. 1997; Staudt et al. 2002; Wu et al. 2015; Haberstroh et al. 2018; Kreuzwieser et al. 2021; Daber et al. 2025). Beyond their biogenic importance, VOCs are highly reactive in the atmosphere and have a strong impact on atmospheric chemistry and air quality (Atkinson and Arey 2003; Folberth et al. 2006; Guenther et al. 2012). These examples illustrate the need for continuous measurements of leaf gas exchange and VOC emissions in forested ecosystems, which remain scarce.

Moreover, leaf-level fluxes strongly impact stem-level fluxes (Andrade et al. 1998; Köcher et al. 2009), however, only few studies have assessed the seasonal and diurnal coordination between the two levels so far (Peters et al. 2023; Peters et al. 2025). Various studies demonstrated a time lag of up to two hours between sap flow rates at the stem top and at the stem base, which they explained by the withdrawal of stem water reserves contributing 9-25% to total daily water use (Phillips et al. 2003; Cermak et al. 2007; Köcher et al. 2013; Leuschner et al. 2024). High stem hydraulic capacitance can significantly enhance drought-tolerance of species and reduce daily fluctuations of water potentials (Betsch et al. 2011; Leuschner et al. 2024). Depletion of stored stem water due to soil water limitations results in decreased predawn water potentials ultimately triggering stomatal closure (Peters et al. 2025). Moreover, stomata sensitively respond to high vapor pressure deficit (*VPD*), which is predicted to increase in the future (Novick et al. 2016; Grossiord 2020; Werner et al. 2025), but due to microclimatic buffering inside the canopy, this response will differ between the sun and shade crown (Niinemets 2023).

Microclimatic gradients inside the tree crown are driven by vertical gradients of photosynthetic photon flux density (*PPFD*), resulting in decreased air temperature (*T_air_)* and *VPD* in shaded areas of the canopy compared to the surrounding atmosphere (Dai et al. 2004; Zellweger et al. 2020; De Frenne et al. 2021). Consequently, shade foliage responds less pronounced to edaphic drying, since concurrent atmospheric water demand and the need for stomatal regulation are lower (Niinemets et al. 2004b; Peltoniemi et al. 2012; Richter et al. 2022). Contrarily, sunlit foliage is more prone to water limitations and therefore usually operates with a higher water use efficiency and earlier stomatal closure than the shade canopy (Niinemets et al. 2004b; Valladares et al. 2016). Vertical microclimatic buffering highly depends on air mass exchange (Flerchinger et al. 2015), canopy density (De Frenne et al. 2021; Gillerot et al. 2021; Zhang et al. 2022), tree species composition and their respective shade casting ability (Ehbrecht et al. 2019; Zellweger et al. 2019; Wang et al. 2025). Microclimatic buffering capacity within the canopy is assumed to mitigate negative impacts of global warming on forest ecosystems (He et al. 2018; De Frenne et al. 2019), and might even enhance the current contribution of the shade canopy to gross primary productivity of 50% (He et al. 2018) to 70% (Sprintsin et al. 2012).

Contributions of shade foliage to whole-canopy carbon gain is determined by *PPFD* penetration into the canopy. High foliage clumping, as in coniferous species, reduces *PPFD* penetration, but at the same time enhances penumbra of foliage, i.e. the partial diffuse shade from the shadow cast of other leaves, which redistributes light more evenly within the crown (Stenberg 1998; Miyashita et al. 2012; Way and Pearcy 2012). Generally, augmenting the diffuse fraction of incoming light, e.g. by aerosol scattering or cloud coverage, enhances whole-canopy carbon gain due to increased carbon assimilation of shade foliage (Dai et al. 2004; Knohl and Baldocchi 2008; Zhou et al. 2021), since diffuse light better penetrates into the canopy and reduces the total area of fully shaded leaves (Roderick et al. 2001; Gu et al. 2002; Urban et al. 2007). Higher light availability in the shade under overcast conditions can also increase VOC emissions of forest stands due to higher contributions of the shade foliage (Laffineur et al. 2013).

Since both leaf gas exchange and VOC emissions are controlled by microclimatic gradients, simultaneous in-situ measurements in various canopy layers are crucial to better understand responses of forest canopies to meteorological conditions and validate results of current modelling approaches (e.g. Niinemets et al. 2002b, Sprintsin et al. 2012, Guenther et al. 2012, He et al. 2018, Chang et al. 2018). Due to logistic challenges, there are only few studies measuring leaf-level gas exchange and VOC emissions continuously in tall tree canopies (Hakola et al. 2006; Kolari et al. 2007; Kolari et al. 2009; Aalto et al. 2014; Werner et al. 2021). Most data on gas exchange and VOC emissions in mature stands were derived from campaign-based measurements neglecting either vertical or seasonal and day-to-day variations (e.g. Bertin et al. 1997, Kesselmeier et al. 1997, Staudt et al. 1997, Niinemets et al. 2002a, Pressley et al. 2004, Plaza et al. 2005, Holzke et al. 2006, Niinemets et al. 2010, Šimpraga et al. 2011, Van Meeningen et al. 2016). While diurnal and seasonal dynamics of total ecosystem net carbon, water and VOC exchange can be estimated by eddy covariance measurements (Laffineur et al. 2013; Haberstroh et al. 2022; Pohl et al. 2023; Scapucci et al. 2024), we lack continuous data on real-time regulation processes of VOC emissions, corresponding leaf gas exchange and the coordination between stem- and leaf-level fluxes, which can significantly enhance our understanding of processes driving seasonal dynamics under varying meteorological conditions.

To this end, we developed and installed a continuous leaf gas exchange measurement system in a tall tree canopy to assess seasonal dynamics of sun and shade foliage. We sampled VOC emissions in regular campaigns from July to October and measured sap flux density, tree water deficit and radial growth in the tree stems to gain a holistic picture of whole-tree carbon and water fluxes. We selected a forest stand containing a deciduous (*Fagus sylvatica*) and a coniferous species (*Pseudotsuga menziesii*) which differed in canopy architecture, foliage clumping (Stenberg 1998; Miyashita et al. 2012), water use strategy (Schumann et al. 2024; Paligi et al. 2025) and VOC storage capacity (Lerdau et al. 1995; Lerdau et al. 1997; Holzke et al. 2006; Holzke et al. 2006) to answer the following research questions: (1) Are there seasonal differences of leaf gas exchange and VOC emissions between both species? (2) How do stomatal regulation and VOC emissions differ between the sun and shade canopy? (3) How are stem-level and leaf-level water fluxes coordinated during the day? Thereby, we aim to better understand seasonal and within-canopy dynamics of leaf gas exchange and VOC emissions resulting of changing *PPFD*, *VPD* and *T_air_.* We further wanted to gain new insights into the relationship between diurnal stem and leaf-level water fluxes regulation.

## Materials and Methods

### Field site and experimental design

The field site is located in the Black Forest on a plateau at an elevation of 520 m a.s.l. close to Ettenheim, Baden-Württemberg, Germany (Tesch et al. 2025) and part of the ECOSENSE collaborative research centre (Werner et al. 2024). Soil in this area is classified as a carbonate-free Cambisol with a silty loam to loamy clay texture originating from sandstone (Werner et al. 2024). Long-term mean *T_air_* and mean annual precipitation of the region was 11°C and 911 mm, respectively, from 1991 to 2020 (Tesch et al. 2025). The stand is mainly composed of European beech (*Fagus sylvatica* L.), but also contains Norway spruce (*Picea abies* (L.) H. Karst.), Silver fir (*Abies alba* Mill.), European larch (*Larix decidua* Mill.), Sessile oak (*Quercus petraea* (Matt.) Liebl.), and larger patches of Douglas fir (*Pseudotsuga menziesii* (Mirbel) Franco). The site is equipped with three canopy access towers, one in a patch dominated by *F. sylvatica*, one in a patch dominated by *P. menziesii* and one in a more mixed patch, where both species interact. Five individuals of *F. sylvatica* and five individuals of *P. menziesii* were selected around a canopy-access tower which provided access to the sun (26 m) and shade (24 m) crowns of three individuals of each species. Diameters at breast height of these individuals ranged between 50.1±11.2 cm and 37.4±5.5 cm for *P. menziesii* and *F. sylvatica*, respectively. Tree height and age varied between 27-30 m and 45-50 years for *P. menziesii* and 24-28 m and 55-110 years for *F. sylvatica*.

### Environmental data

In February 2023, a meteorological station was installed on an open area 250 m southwest of the field site. The station continuously measured shortwave radiation using a CNR4 sensor (Kipp & Zonen, Delft, Netherlands), *T_air_* and relative humidity (*RH*) with a HygroVUE5 sensor (Campbell Scientific Ltd., Shepshed, UK) as well as precipitation with a Young 52202 Tipping Bucket (R.M. Young Company, Traverse City, MI, USA). Additionally, in Mai 2024, a LI-190 quantum sensor (LI-COR Environmental, Lincoln, NE, USA) was installed above the canopy (46 m height) to measure *PPFD* and two HygroVUE10 sensors (Campbell Scientific Ltd., Shepshed, UK) were in installed in 27 m and 18 m height to measure *T_air_* and *RH* in the sun and shade canopy, respectively. Data were recorded on CR1000 data loggers (Campbell Scientific Ltd., Shepshed, UK) every minute. *VPD* was calculated from *T_air_* and *RH* using the guideline provided by the WMO (WMO 2018). *PPFD* at the open area was calculated from shortwave radiation data using a linear regression model with the *PPFD* sensor above the canopy.

At four locations four SMT100 soil moisture sensors (Truebner GmbH, Neustadt, Germany) were installed at four depths (0.30 m, 0.50 m, 0.70 m, 0.90 m). Additionally, 35 SMT100 sensors were distributed radially around the measurement trees at 0.05 m depth. Sensors recorded volumetric soil water content (*VWC*) every 15 minutes on a data logger (CR350, Campbell Scientific Ltd., Shepshed, UK). Data quality of *VWC* was checked using the quality control procedure by Dorigo et al. (2013) and only values with a ‘good’ quality flag were selected for analysis.

Leaf water potentials (*ψ_leaf_)* were measured once a month on small branchlets of the sun crown of all individuals which could be reached by the canopy-access tower using a Scholander pressure chamber (Soil Moisture Equipment, Santa Barbara, CA, USA, Scholander et al. 1965). Predawn water potential (*ψ_PD_*) was measured in the two hours prior to sunrise and midday water potential *(ψ_MD_)* in the two hours around solar noon from May to October 2024.

### Sap flow and dendrometer measurements

In autumn 2023, tree stems were equipped with heat pulse velocity sensors (SFM5, UGT, Muencheberg, Germany) at breast height (1.3 m above the ground) to determine sap flux density (*J_s_*) at two different xylem depths (10 and 20 mm < the cambium). Insulating foil was wrapped around the sensor heads to protect them from UV-radiation and *T_air_* fluctuations. Measured data were recorded on a data logger (CR1000, Campbell Scientific Ltd., Shepshed, UK) every 15 minutes and transmitted in real-time to the ECOSENSE data base (Tesch et al. 2025).

The Dual Method Approach (Forster 2019, 2020), combining the heat ratio method (HRM) (Burgess et al. 2001; Marshall 1958) and the Tmax method (Cohen et al. 1981), was used to calculate *J_s_* separately for the inner and outer xylem. Total *J_s_* of the active sapwood was then calculated by scaling *J_s_* of inner and outer xylem to the respective proportion of the sapwood area measured by the thermistors. For the determination of sapwood area, tree cores were drilled in October 2024 at breast height and illuminated immediately. Translucent zones were classified as functional sapwood with active water transport (Munster-Swendsen 1987; Quiñonez-Piñón and Valeo 2018). Active sapwood area was 158.8 ± 35.3 cm^2^ in *P. menziesii* and 117.6 ± 17.4 cm^2^ in *F. sylvatica*. Needle probe misalignment was corrected by solving HRM and Tmax equations for needle distances (Kinzinger et al. 2024; Dumberger et al. 2025). During nights (*PPFD* < 10 µmol m^-2^ s^-1^) with *VPD* < 0.1 kPa, no sap flow was expected, resulting in heat velocity values equal to zero in both equations. Corrected needle distances between the 5% and 95% percentiles of each individual tree were then interpolated for all nights with non-zero flow conditions. Two sensors in two individuals of *F. sylvatica* failed temporarily leading to data gaps. Linear regression models with an adjacent tree, which showed a similar trend of *J_s_* (Spearmańs rho: 0.81-0.91), were calculated and used to fill the data gaps.

Custom-made point dendrometers as described in Wang and Sammis (2008) were installed on the stems at 1.3 m above the ground to record stem radius variation as in Dumberger et al. (2025). A potentiometer (Model 9605 BEI, Duncan Electronics, Commerce, Texas, USA) mounted to a metal angle bracket and two stainless-steel rods was attached to the trees directly touching the stem (Wang and Sammis 2008). In all individuals of *P. menziesii* the upper layers of bark were removed gently to ensure direct contact to water conducting wood tissue. Sensors were connected to CR1000 data loggers (Campbell Scientific Ltd., Shepshed, UK) and recorded data at an interval of 15 minutes. Raw data were further processed with the “treenetproc” package which was used to remove outliers and calculate radial growth and tree water deficit (*TWD*) by the zero-growth approach (Zweifel et al. 2016, Knüsel et al. 2021). During the whole vegetation period of 2024, values of one *F. sylvatica* individual had to be removed from the data set due to sensor failure.

### Leaf and needle cuvettes

For the measurements of leaf gas exchange and VOC emissions, two different types of leaf enclosure systems were used. Single leaves of *F. sylvatica* were equipped with a novel, lightweight and minimally invasive cuvette (“ECOvette”, Frey et al. 2025a, Figure 1, Supplementary Material Figure S1 A & B). The ECOvette comprises a hemispherical capsule which is attached to the leaf by two ring magnets with an ultra-soft silicone seal and covers a leaf area of 6.16 cm^2^. ECOvettes were equipped with an *T_air_* and *RH* sensor (SHT40, Sensirion GmbH, Gerlingen, Germany) inside and outside of the cuvette and a thermocouple (P TF-50, Type K, PeakTech Prüf- und Messtechnik GmbH, Ahrensburg, Germany) to measure leaf temperature (*T_leaf_*, Frey et al. 2025b). Sensor values were sent every ten seconds via a Wifi from a microcontroller (ESP8266 Wemos D1 Mini, Espressif Systems, Shanghai, China) to a Raspberry Pi (Raspberry Pi Trading Ltd., Cambridge, UK) which logged the data on a central database (Tesch et al. 2025). A more detailed description and a proof of concept for gas measurements with the ECOvette can be found in Frey et al. (2025a).

**Figure 1:**
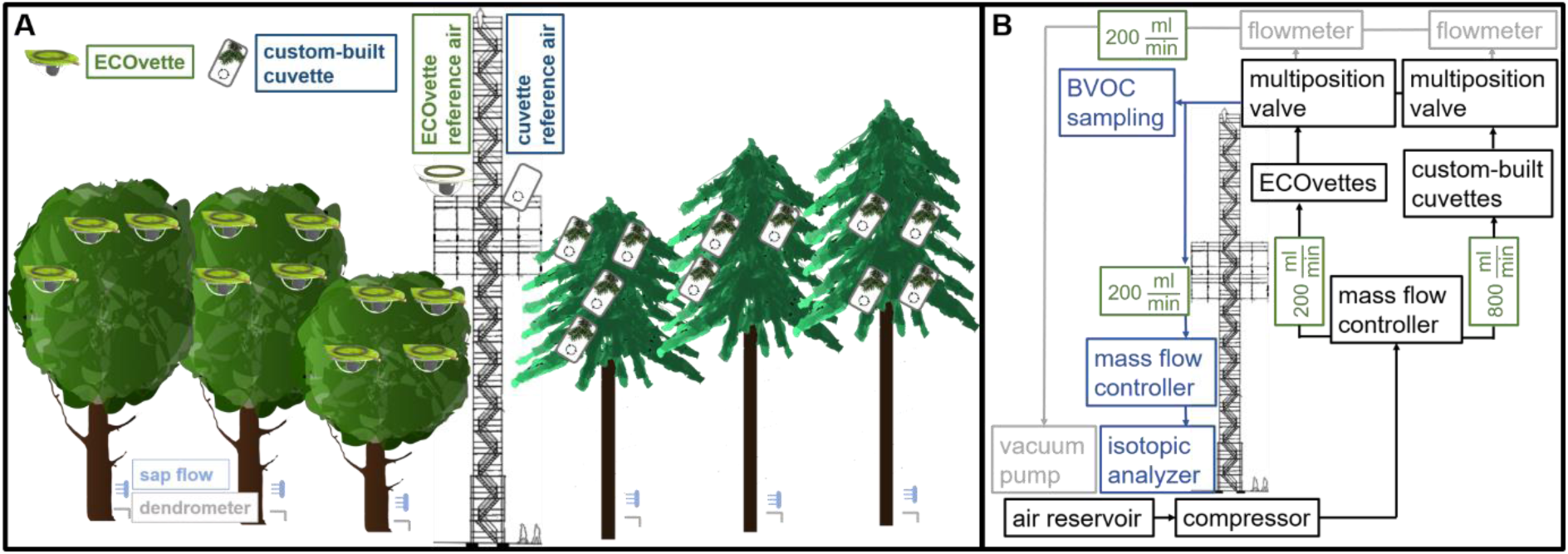
Schematic overview of the automated measurement system. The left panel (A) shows the distribution of sun and shade cuvettes in the canopy of *F. sylvatica* (n=3) and *P. menziesii* (n=3). Empty cuvettes represent the cuvettes used for measuring reference air. The right panel (B) shows the connection between the air supply unit (black color), the measurement unit (blue color) and the constant flushing unit (grey color) of the automated measurement system. Flow rates are indicated in green color.

Small branchlets of *P. menziesii* were enclosed into larger cuvettes custom-built for conifer branchlets (Werner et al. 2021) made of FEP film (PTFE-Spezialvertrieb GmbH, Stuhr, Germany) and a supporting frame of PFA-tubing (1/4”, Wolf-Technik eK, Stuttgart, Germany). Cuvettes were equipped with small fans (MF40101V2-1000-UA99, Sunon, Kaohsiung City, Taiwan) to ensure homogenous mixing of the air and sealed air-tight with teroson (Teroson RBII, Henkel AG & CoKG, Düsseldorf, Germany) and rubber band (Supplementary Material Figure S1 C). We additionally attached the environmental sensors of six ECOvettes to three conifer cuvettes in the sun and shade canopy, respectively, to measure *T_air_*, *T_leaf_* and *RH*. At the end of the measurement period or if cuvettes were damaged after thunderstorms, branchlets were harvested and projected needle area was determined in the laboratory using a commercial scanner connected to the GSA Image Analyzer Software (GSA GmbH, Rostock, Germany). Needles were detached from the branchlets and spread evenly on the scanner surface with as minimum overlapping as possible.

Twelve ECOvettes and twelve conifer cuvettes were installed in the sun (n=6 per species) and shade (n=5 per species) canopy of three individuals of *F. sylvatica* and *P. menziesii*, respectively (Figure 1A). We measured *PPFD* at the cuvettes with a hand-held light meter (LI-250A, LI-COR Environmental, Lincoln, NE, USA) on several days throughout the entire study period to ensure proper shading of the shade cuvettes. Shade foliage received on average 9% (53.7 ± 24.7 µmol m^-2^ s^-1^) and 8% (33.8 ± 3.8 µmol m^-2^ s^-1^) of *PPFD* received by sun foliage of *F. sylvatica* (555.9 ± 71.9 µmol m^-2^ s^-1^) and *P. menziesii* (544.9 ± 111.4 µmol m^-2^ s^-1^), respectively. One cuvette of each type was left empty as a reference for the incoming air into the cuvettes. A small piece of FEP film (PTFE-Spezialvertrieb GmbH, Stuhr, Germany) was used to seal the empty ECOvette gas-tight.

Environmental sensors of the ECOvettes failed during several periods due to technical issues. Data gaps were filled using linear regression models with adjacent sensors (R^2^ = 0.68-0.99), if available, or with values measured at the canopy-access tower (R^2^ = 0.47-0.98) in 27 m height (sun leaves) and 18 m height (shade leaves). For six conifer cuvettes, which were not equipped with environmental sensing, *T_air_*, *T_leaf_* and *RH* values from adjacent sun and shade cuvettes were used. Mounting of the thermocouples to the needles of *P. menziesii* was challenging, and constant direct contact with needles could not be ensured. We thus used the temperature inside the cuvette (*T_cuv_*) for calculations of gas exchange and further analysis of microclimatic conditions.

### Automated air sampling systems

An automated air sampling system (Figure 1B) was established in the forest following the approach already used in climate chambers (Fasbender et al. 2018; Werner et al. 2020) and in an artificial tropical rain forest (Werner et al. 2021).

Air was supplied to the cuvettes from the tower base by a compressor (AG OFCAS 36 Plus, LNI Swissgas GmbH, Kamen, Germany) connected to a buffer tank with a volume of 100 L (LNI Swissgas GmbH, Kamen, Germany), which sent compressed air to the tower platform (26 m height) through PFA-tubing (1/4”, Wolf-Technik eK, Stuttgart, Germany) with a constant pressure of 2 bar. On the tower platform, the supplied air was distributed to 24 thermal mass flow controllers (G-Series, Brooks Instrument GmbH, Dresden, Germany) by two custom aluminum manifolds. Mass flow controllers regulated the air flow to the ECOvettes to 350 ml min^-1^ and to the conifer cuvettes to 800 ml min^-1^ accounting for the differing air volume of the cuvettes (5.8 cm^3^ & ∼1000 cm^3^). Supplied air was delivered to the conifer cuvettes by 1/4” PFA tubing and to the ECOvettes by 1/8” PFA-tubing. Sampling air was pumped with a vacuum pump (KNF LABOPORT® N840G, Faust Lab Science GmbH, Klettgau, Germany) situated at the tower base to a distribution unit at the tower platform consisting of two flow-through multiposition valves (16 positions, 1/8”, Valco Instruments Co. Inc., Schenken, Switzerland) which automatically switched between the 24 cuvettes. Sampled air from the measured cuvette was sent through 1/4” PFA tubing at a rate of 200 ml min^-1^ controlled by a thermal mass flow controller (G-Series, Brooks Instrument GmbH, Dresden, Germany) to an Isotope and Gas Concentration Analyzer (G-2131-i, Picarro Inc., Santa Clara, USA) at the tower base measuring ^12^CO_2_, ^13^CO_2_ and H_2_O concentrations every two seconds. Remaining cuvettes were constantly flushed at a flow rate of 200 ml min^-1^ controlled by flowmeters (SHLLJ, Sorekarin, Yueqing, China) to ensure continuous supply of fresh CO_2_ and avoid condensation of transpired H_2_O. Switching between the cuvettes was realized every nine minutes, of which the last 60 seconds were averaged. For data quality control, the standard deviation of the averaged 60 s interval was calculated and measurements exceeding a standard deviation of 2.5 ppm (^12^CO_2_) and/or 0.05 % (H_2_O) were removed from the data, as the measurement was likely not stable. Additionally, 16 full days and several single data points were removed due to maintenance work at the cuvettes, damage of the cuvettes or failure of the whole measurement system. After this rigid data cleaning process, 65% of the initial data (6452 of 9964 data points) were further analyzed.

### Calculation of leaf gas exchange

For the calculation of leaf gas exchange, ^12^CO_2_, ^13^CO_2_ and H_2_O concentrations of the air entering the leaf cuvettes were assumed to be equal to the air measured inside the reference air cuvettes. Reference air cuvettes were measured every 54 minutes and extended to the entire data set by linear interpolation. Differences between ^12^CO_2_, ^13^CO_2_ and H_2_O concentrations of the interpolated reference air and the air exiting the leaf cuvettes were then used to calculate transpiration (*E,* Supplementary Material Formula S1), net carbon assimilation (*A_net_,* Supplementary Material Formula S2), stomatal conductance for water vapor (*g_s_*, Supplementary Material Formula S3), leaf discrimination for ^13^C during photosynthesis (*Δ_leaf_,* Supplementary Material Formula S4), water use efficiency (*WUE*, Supplementary Material Formula S5) and intrinsic water use efficiency (*WUEi,* Supplementary Material Formula S6) of enclosed leaves and branchlets based on von Caemmerer and Farquhar (1981) and Evans et al. (1986).

### Measurements of biogenic volatile compound emissions

For the measurements of VOC emissions, an additional 1/4” PFA tee connector and a short piece of 1/4” PFA tubing were installed between the PFA tubing coming from the cuvettes and the two multi-position valves (Figure 1B). On thirteen days from July to October, VOC emissions were collected from the cuvettes between 11 am to 2 pm on glass tubes filled with Tenax (Sigma-Aldrich, Munich, Germany) at a rate of 50 ml min^-1^ for one hour with pumps (Pocket Pump Touch, SKC, Dorset, UK). After sampling, tubes were stored in airtight vials (Labco Limited, Lampeter Ceredigion, United Kingdom) in the fridge (4°C) until they were measured with a Gas Chromatograph (GC, Model 7820A and 7890B, Agilent Technologies, Santa Clara, USA), see Kreuzwieser et al. (2021) for a detailed description. Briefly, samples were heated to 240°C in a thermal desorption unit (TDU, Gerstel, Mühlheim a.d. Ruhr, Germany), thereafter cryotrapped at -70°C (KAS, Gerstel, Mühlheim a.d. Ruhr, Germany), reheated to 240°C at a rate of 12°C s^-1^ and subsequently released into a separation column (DB-5MS UI, Agilent Technologies, Santa Clara, USA). For the separation of the compounds the column was heated for 54 minutes in three steps from 45°C to 280°C. Separated compounds were then introduced into the mass spectrometer, operating at 70.35 eV, an ion source temperature of 230°C and a quadrupole temperature of 150°C. Measured mass spectra were analyzed using the MassHunter Software (Agilent Technologies, Santa Clara, USA). Concentrations of the compounds were quantified using a standard mixture composed of eight compounds (isoprene, ⍺-pinene, β-pinene, limonene, sabinene, trans-β-ocimene, caryophyllene and farnesene). VOC fluxes were calculated using Formula S7 (Supplementary Material).

### Data analysis

Data analysis was performed using the software R (version 4.5.1) and RStudio (version 2025.09.1, R Core Team 2025)).

For the analysis of seasonal dynamics and diurnal courses we classified the vegetation period into three periods. We selected these three periods since *T_air_*, *VWC* at 0.05 m depth and *VPD* differed, indicative for a potential heat and/or drought stress. Meteorological conditions during early summer (18/06/2024 to 19/07/2024) were warm and wet, during midsummer (24/07/2024 to 29/08/2024) we recorded the warmest period with highest *VPD* and lowest *VWC*, and during late summer (09/09/2024 to 18/09/2024) cold conditions with low *VWC* were observed (Table 1). Furthermore, continuous data from our leaf gas exchange measurements system were available during all three periods.

**Table 1:** Daily minimum soil water content (*VWC)* in 0.05 m depth, daily maximum vapor pressure deficit (*VPD*) and daily maximum air temperature (*T_air_*) ± the standard error for the respective variable averaged over the three analyzed periods.

|  | early summer<br>(18<sup>th</sup> Jun - 19<sup>th</sup> Jul) | midsummer<br>(24<sup>th</sup> Jul - 29<sup>th</sup> Aug) | late summer<br>(9<sup>th</sup> Sep – 18<sup>th</sup> Sep) |
| --- | --- | --- | --- |
| **VWC [Vol. %]** | 26.6 ± 0.3 | 17.1 ± 0.4 | 16.9 ± 0.2 |
| **VPD [kPa]** | 1.2 ± 0.1 | 1.5 ± 0.1 | 0.4 ± 0.1 |
| **T<sub>air</sub> [°C]** | 23.8 ± 0.7 | 26.2 ± 0.6 | 14.1 ± 0.5 |

We computed three different linear mixed effect models (package “lme4”) using the three tree individuals of each species as a random effect to account for repeated measurements (Bates et al. 2015). We further analyzed the models comparing contrasts of estimated marginal means (package “emmeans”) for the respective predictor variables (Lenth 2023). Model assumptions were checked with the package “performance” and, if necessary, response variables were transformed as specified below (Lüdecke et al. 2021). The first model calculated differences of *J_s_*, radial growth and *TWD* between the two species on a monthly resolution. A cube root transformation was applied to *J_s_* and a square root transformation to radial growth and *TWD*, respectively, to ensure normality of residuals and homogeneity of variance. The second model identified differences between the measurement campaigns and between the species for predawn and midday *ψ_leaf_*. The third model analyzed differences between log-transformed VOC emission rates of the sun and shade foliage for each of the thirteen sampling dates. Additionally, to the estimated marginal means a type III ANOVA was used as a post-hoc test to analyze seasonal effects influencing VOC emissions.

For the analysis of diurnal courses, two different generalized additive models (package “mcgv”) were fitted and estimated marginal means of the models were compared (Wood 2017; Lenth 2023). The first model was fitted for *T_cuv_*, *VPD*, *g_s_*, *A_net_* and *E* with species, sun/shade exposition and the three predefined periods as predictors, a smoothing spline for the interaction of the three predictors, a thin-plate regression spline for time of a day on a half-hourly resolution and tree individual as a random effect. The second model was fitted for normalized values of J_s_, *g_s_*, *E*, *VPD* and *TWD* with species and the three predefined periods as predictors, a smoothing spline for the interaction of the two predictors, a thin-plate regression spline for time of a day on a 15-minute resolution and tree individual as a random effect. The second model was fitted separately for the sun and shade foliage. Normalized values were obtained for each tree individually by dividing the respective variable by the 99% quantile of maximum values over the whole season.

## Results

### Meteorological conditions at the field site

The summer of 2024 was dominated by wet and warm conditions with no pronounced heat waves, but two cold spells (beginning of July and mid-September) with a sudden drop of 10 K in *T_air_* (Figure 2 A&B). *VWC* remained high in all layers during early summer, while a slight dry-down period was recorded from mid-July to early September, when *VWC* in 0.05 m depth dropped to 12.9 ± 1.4 % (Figure 2C). *J_s_* resembled meteorological conditions with pronounced reductions during the colder periods (Figure 2D). *J_s_* of *F. sylvatica* tended to be higher than *J_s_* of *P. menziesii* during July and August. During the other months, *J_s_* of *P. menziesii* slightly exceeded *J_s_* of *F. sylvatica*, albeit this difference was only significant during March and April (p < 0.01), when leaves of *F. sylvatica* were not yet fully developed. Radial growth started earlier in coniferous *P. menziesii*, but daily growth rate was only significantly higher compared to *F. sylvatica* during April (p < 0.05). There was no difference in the daily growth rate between the species during May, but during June and July, *F. sylvatica* exceeded the daily growth rate of *P. menziesii* (p = 0.05) which resulted in a slightly higher total radial increment for *F. sylvatica* (3.6 ± 0.7 mm) than for *P. menziesii* (3.2 ± 0.9 mm, Figure 2E). A significant *TWD* was absent for both species until the end of June; however, coinciding with decreasing *VWC* (Figure 2C), *TWD_min_* of *P. menziesii* increased in August to 35.5 ± 33.8 µm and stayed elevated until October. For *F. sylvatica*, *TWD_min_* only reached maximum of 12.1 ± 6.0 µm at the end of the study period in October (Figure 2F).

**Figure 2:**
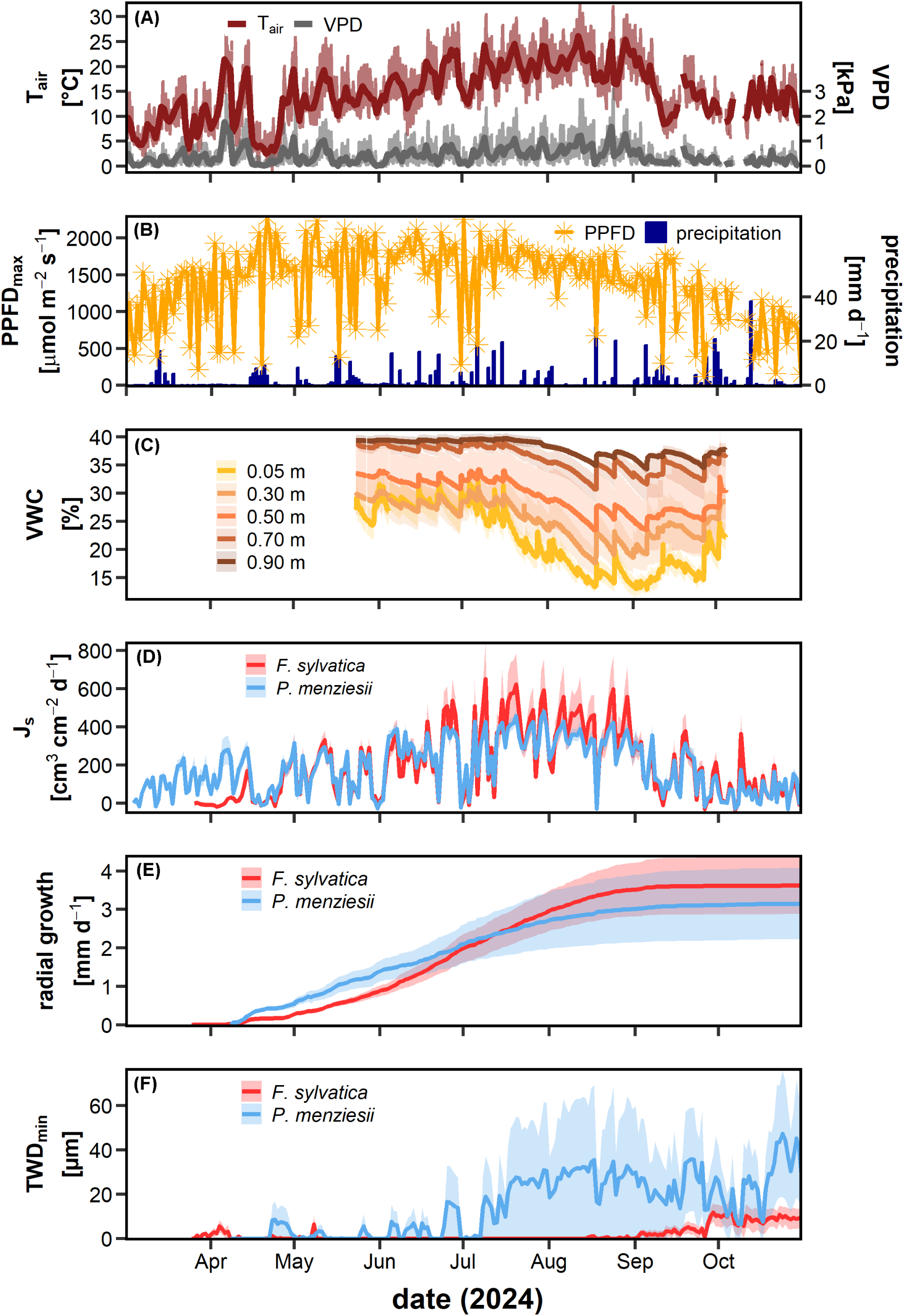
Environmental conditions (A-C) as well as stem-level water and carbon fluxes (D-F) at the field site during the vegetation period of 2024. Panels A-C show (A) daily mean air temperature (*T_air_*, dark red line) and *T_air_* on a 5-minute resolution (bright red line), daily mean vapor pressure deficit (*VPD*, dark grey line) and *VPD* on a 5-minute resolution (bright grey line), (B) daily maximum photosynthetic photon flux density above the canopy (*PPFD_max_*, yellow points and line) and daily sum of precipitation (blue bars), and (C) volumetric soil water content (*VWC*) in five depths (0.05, 0.30, 0.50. 0.70. 0.90 m). Panels D-F show (D) daily sum of sap flux density (*J_s_*), (E) cumulative radial growth, and (F) daily minimum tree water deficit (*TWD_min_*) of *F. sylvatica* (red line, n=5) and *P. menziesii* (blue line, n=5), respectively. Shaded areas in panel C-F show the standard error, while lines show the mean values of four soil pits (C) and five tree individuals (D-F), respectively. Due to sensor failure after heavy rainfalls during spring and autumn, available data of *VWC* (C) were limited to the period between June and October.

### Seasonal leaf gas exchange

Leaf gas exchange differed substantially not only between the two investigated species, between sun and shade foliage and between predefined periods (Figure 3). On average, daily maximum *A_net_* and *E* measured at single leaves of *F. sylvatica* were higher than rates measured at small branchlets of *P. menziesii* (Figure 3 C-F) and showed more pronounced fluctuations in response to changing weather conditions. However, comparisons between the two species need to be interpreted carefully, since penumbral effects in measured shoots of *P. menziesii* cannot be excluded.

**Figure 3:**
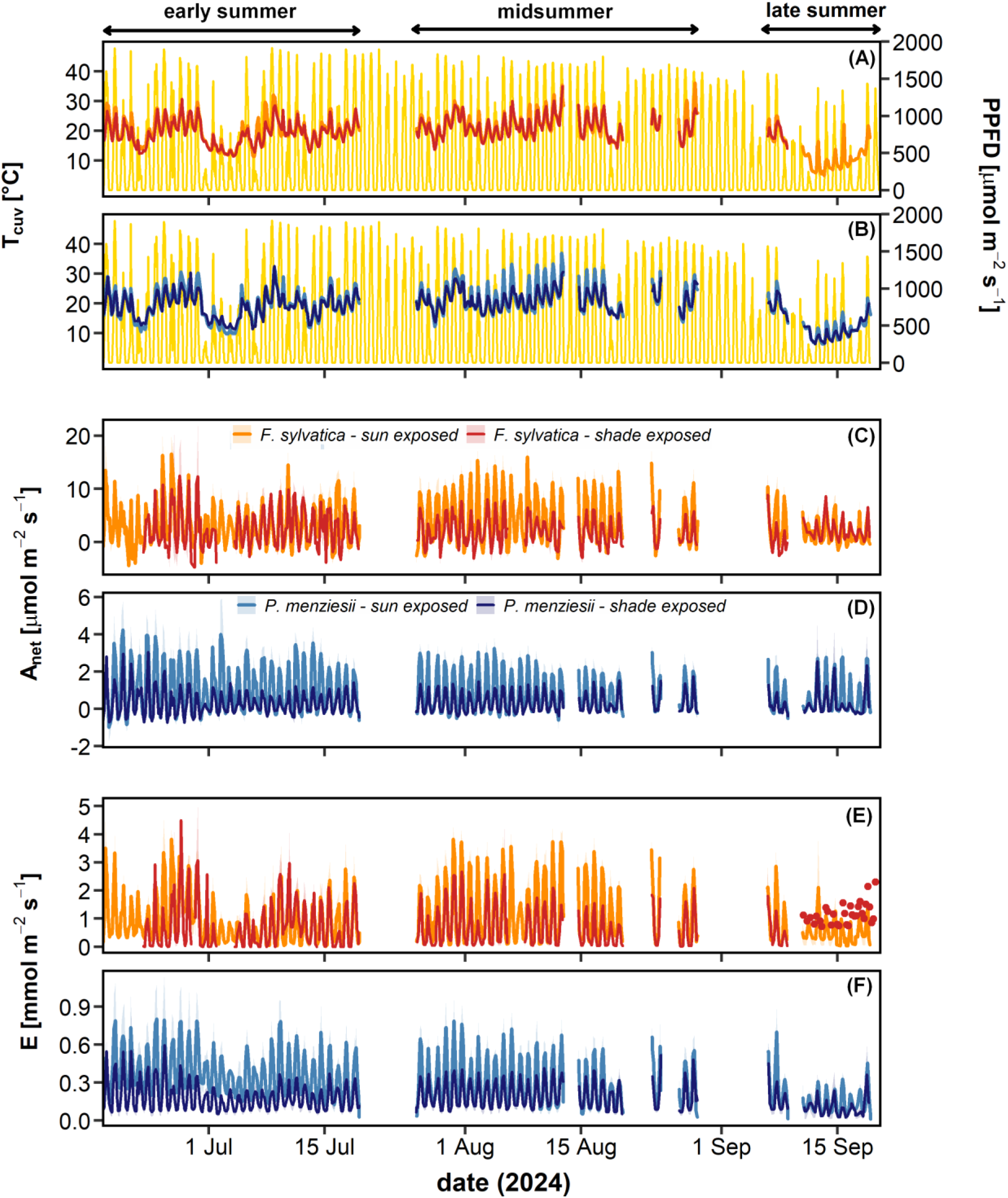
Continuous measurements of carbon and water fluxes on the leaf-level from June to September 2024. Lines and shaded areas display the mean and standard error of all measured cuvettes (n=5-6) on an hourly resolution, respectively. Shown are (A & B) the temperature inside the cuvette (*T_cuv_*) and photosynthetic photon flux density measured at the top of the tower in 46 m height (*PPFD*, yellow line), (C & D) net carbon assimilation rate (*A_net_*), and (E & F) the transpiration rate (*E)* of single leaves of *F. sylvatica* (A, C, E, n=3) in the sun (orange line) and shade (red lines) canopy and of small branchlets of *P. menziesii* (B, D, F, n=3) in the sun (bright blue line) and shade (dark blue line) canopy. Large gaps were caused by technical failure of the measurement system due to heavy wind, rain or heat. Due to low data availability, transpiration of shade leaves of *F. sylvatica* in panel E is displayed as single points. Note the different y-axis scales between panel C and D and between panel E and F.

Differences between the predefined periods were observed in the comparison of shade and sun foliage (mean daily maxima in brackets): during early summer, *A_net_* of shade leaves of *F. sylvatica* (9.8 ± 0.7 µmol m^-2^ s^-1^) nearly reached the same rates as in sun leaves (11.1 ± 0.5 µmol m^-2^ s^-1^), whereas during midsummer *A_net_* of sun exposed leaves (15.5 ± 0.5 µmol m^-2^ s^-1^) clearly exceeded shade exposed leaves (6.7 ± 0.6 µmol m^-2^ s^-1^) and the same pattern was also observed in *E* (Figure 3C&E). A pronounced decrease of *A_net_* and *E* in the sun foliage of *F. sylvatica* was visible during the two periods with low *T_air_* and low *PPFD* at the beginning of July and in mid-September, while rates of the shade foliage remained relatively stable over the season and were less affected by changes in *PPFD* and *T_cuv_*.

*A_net_* of the sun branchlets of *P. menziesii* exceeded rates of the shade branchlets during most of early summer (sun: 3.5 ± 0.2 µmol m^-2^ s^-1^ and shade: 1.6 ± 0.1 µmol m^-2^ s^-1^) and midsummer (sun: 3.1 ± 0.1 µmol m^-2^ s^-1^ and shade: 1.4 ± 0.1 µmol m^-2^ s^-1^), which was also resembled in *E*. Both, *A_net_* and *E*, were generally more influenced by changing meteorological conditions in the sun than in the shade. At the beginning of early summer and especially during the cold spell in September shade foliage of *P. menziesii* reached *A_net_* and *E* rates comparable to the sun foliage.

*Δ_leaf_*, representing the *in-situ* discrimination of the leaf against ^13^C during photosynthesis, fluctuated in response to changing meteorological conditions and showed high diurnal variability (Supplementary Material Figure S2). *Δ_leaf_* of both species showed a pronounced decline by 3-6 ‰ in response to the cold spell in mid-September concurrent with an increase of *WUEi* (Supplementary Material Figure S3). During midsummer, daily mean *Δ_leaf_* was lower in the sun leaves than in the shade leaves of *F. sylvatica* (17.7 ± 0.4 ‰ and 19.8 ± 0.5 ‰, respectively) and *P. menziesii* (21.1 ± 0.4 ‰ and 22.6 ± 0.5 ‰, respectively). A pronounced reduction of *Δ_leaf_* was observed in the second half of midsummer, particularly in *P. menziesii*, which reduced *Δ_leaf_* in the sun leaves by 6 ‰ and in the shade leaves by 7‰.

During this period, we detected a gradual dry-down, when *VWC* at 0.05 m depth decreased to 12.9 ± 1.4 % and *VPD* increased to 3.5 ± 0.1 kPa and 3.4 ± 0.2 kPa in the sun foliage of *F. sylvatica* and *P. menziesii*, respectively (Figure 4 A-C). Concurrently, *ψ_PD_* (-1.0 ± 0.1 MPa) and *ψ_MD_* (-2.2 ± 0.2 MPa) of *P. menziesii* decreased significantly (p < 0.05) compared to the values measured in early summer (*ψ_PD_*: -0.3 ± 0.1 MPa*, ψ_MD_*: -1.2 ± 0.1 MPa) and both were also significantly lower than those of *F. sylvatica* (p < 0.01, Figure 4A). *ψ_PD_* of *F. sylvatica* remained unaffected, while *ψ_MD_* also decreased significantly to -1.7 ± 0.1 MPa compared to early summer (p < 0.001). Sun foliage of *P. menziesii* responded with a gradual decline of *g_s_* during midsummer, reducing maximum *g_s_* by 45% (from 195.2 ± 22.8 to 86.1 ± 12.2 mmol m^-2^ s^-1^), whereas maximum *g_s_* of sun leaves of *F. sylvatica* (188.4 ± 6.5 mmol m^-2^ s^-1^) was unaffected by drier conditions (Figure 4 D&E). Reduced *g_s_* in the sun branchlets of *P. menziesii* was accompanied by a twofold increase of *WUEi* during midsummer compared to early summer, which was not visible in the shade branchlets (Supplementary Material Figure S3). Shade foliage of *P. menziesii* maintained similar *g_s_* throughout the whole study period and reached highest values (88.7 ± 11.3 mmol m^-2^ s^-1^) during midsummer. During midsummer, shade foliage of *F. sylvatica* also slightly reduced *g_s_* by 20% to 85.1 ± 6.2 mmol m^-2^ s^-1^, but recovered again to 101.5 10.9 mmol m^-2^ s^-1^ at the end of the period after a heavy precipitation event.

**Figure 4:**
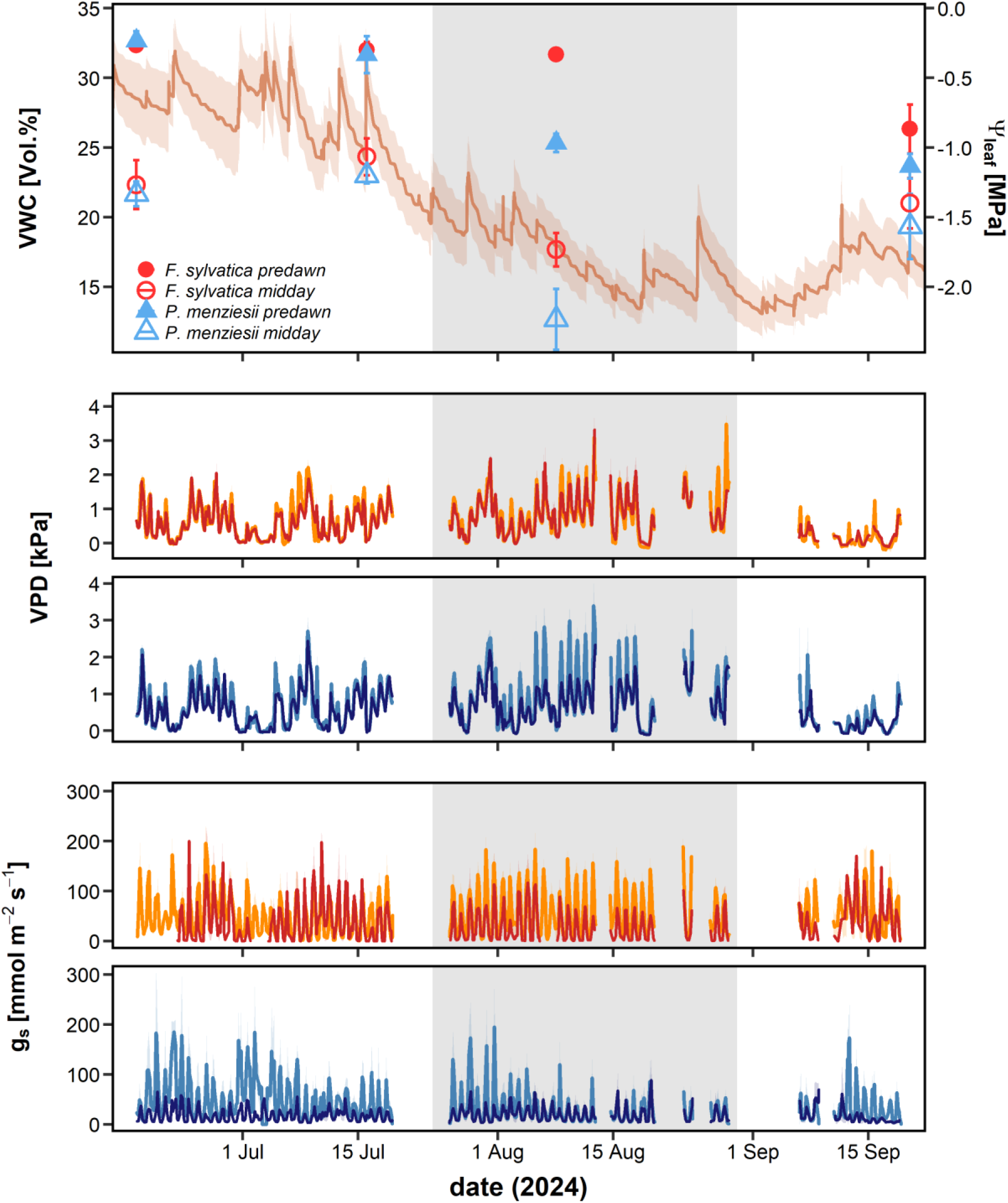
Soil water content in 0.05 m depth (*VWC*, A, brown line), leaf water potentials (*ψ_leaf_,* A) measured on sunlit twigs of *F. sylvatica* (n=3, red points) and *P. menziesii* (n=3, blue triangles) before sunrise (predawn, filled symbols) and around noon (midday, empty symbols), vapor pressure deficit (*VPD)* at the leaf enclosures (B & C) and stomatal conductance (*g_s_*) of enclosed leaves of *F. sylvatica* (D, n=3) and enclosed branchlets of *P. menziesii* (E, n=3) in the sun (bright color) and shade (dark color) canopy. Shaded areas and error bars show the standard error of the respective variable, while lines and points show the mean of all individuals (n=3 per species) and soil pits (n=4). Grey shaded area in the background highlights the period of midsummer.

*Seasonal* VOC *emissions*

For both species, a declining seasonal trend of monoterpene emissions was detected (p < 0.001), which differed between the species (p < 0.001). In sun foliage of *F. sylvatica,* we found a gradual decline of emissions towards late summer. We could not detect any monoterpene emissions by sun leaves of *F. sylvatica* on the last two measurement dates which was probably related to early senescence in the sun canopy. In the sun foliage of *P. menziesii,* two pronounced declines of emissions from early summer to midsummer and from midsummer to late summer were observed (Figure 5 A&B). The first decline occurred simultaneously with decreasing *g_s_* and *ψ_leaf_* during the dry-down period (Figure 4E) and the second during the sudden cold spell (Figure 3B). Monoterpene emissions of the sun foliage of both species were usually higher than those of the shade foliage, albeit this difference was only significant on four days for *F. sylvatica* due to the high variability of emissions between the tree individuals (Figure 5 A&C). On cloudy days, differences between emissions of sun and shade foliage diminished in both species, particularly in *F. sylvatica,* leading to similar monoterpene emissions in both canopy layers.

**Figure 5:**
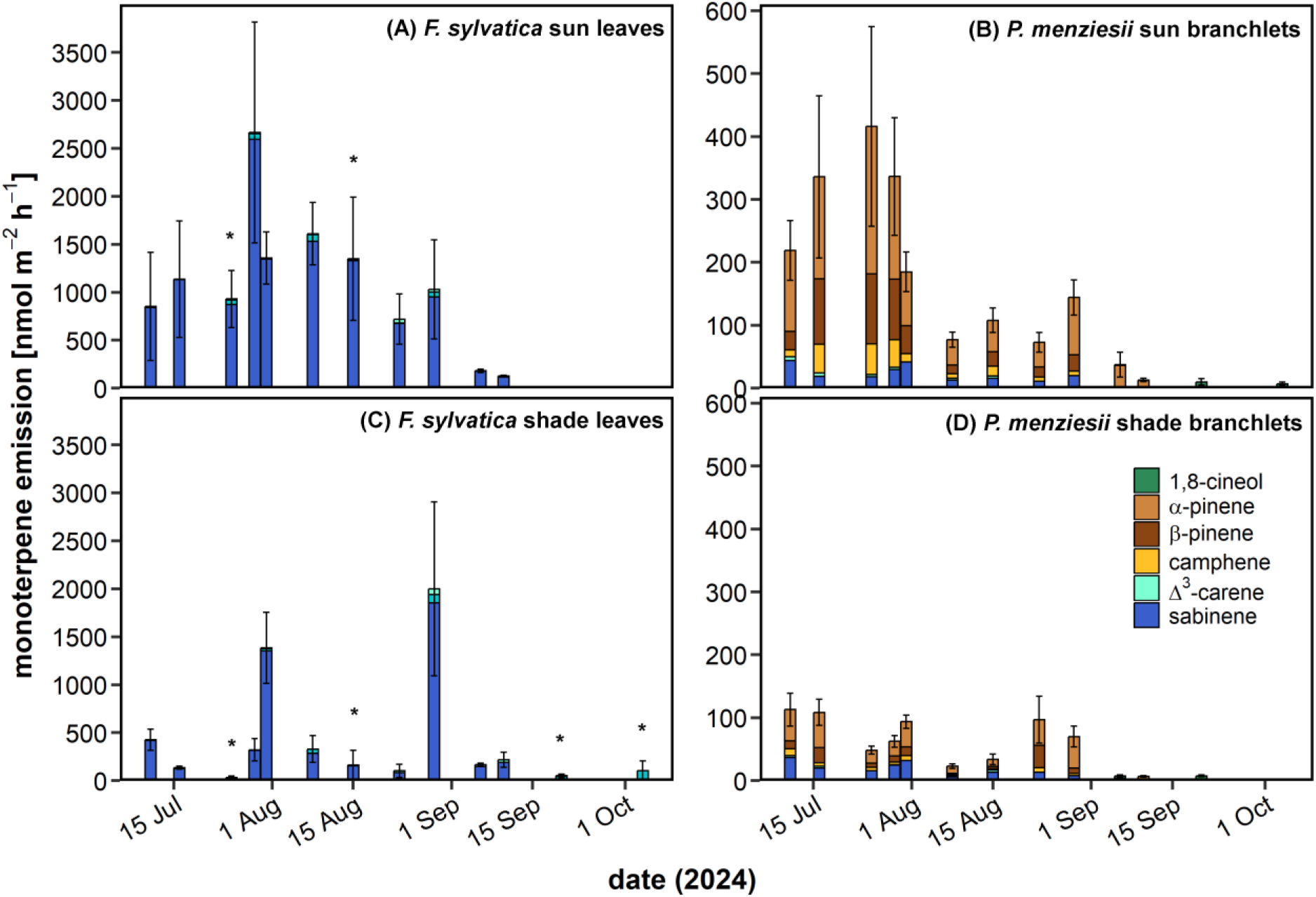
Monoterpene emissions in the sun (A & B) and shade canopy (C & D) of single leaves of *F. sylvatica* (A & C, n=3) and small branchlets of *P. menziesii* (B & D, n=3) measured during 13 campaigns from July to October 2024. Error bars show the standard error calculated with error propagation. Asterisks indicate a significant difference between the monoterpene emission sum of sun and shade leaves (* p < 0.05). Note the different y-axis scale between panels A&B and C&D due to the different emissions rates of the two species.

Overall, 32 compounds could be identified from measured samples of which only the seven monoterpenes were further analyzed. On average, sun leaves of *F. sylvatica* emitted mainly sabinene (89.5 ± 3.3%), but also Δ^3^-carene (9.5 ± 3.4%) and limonene (1.3 ± 0.6%), while in the shade, monoterpenoid emissions were more equally composed of sabinene (52.8 ± 11.4%), limonene (28.3 ± 12.4%) and Δ^3^-carene (23.6 ± 8.9%). *P. menziesii* emitted a more diverse blend of monoterpenoids which was mainly composed of ⍺-pinene (sun: 55.5 ± 5.5%, shade: 42.8 ± 7.1%), sabinene (sun: 20.1 ± 3.8%, shade: 31.7 ± 5.9%) and ꞵ-pinene (sun: 20.1 ± 3.8%, shade: 12.7 ± 1.6%), but also to a lower extent of camphene (sun: 8.5 ±1.3%, shade: 12.1 ± 4.4%) and Δ^3^-carene (sun: 2.7 ± 0.9%, shade: 5.5 ± 3.4%). Additionally, during September and October low emissions of 1,8-cineol (0.6-17.8 nmol m^-2^ s^-1^) were detected in the sun and shade foliage of *P. menziesii*.

### Diurnal courses of leaf gas exchange, sap flux density and tree water deficit

In both species, we found seasonal and within-canopy differences of diurnal courses of leaf gas exchange. During midsummer, microclimatic differences between sun and shade leaves of both species were highest with maximum *T_cuv_* differences of 2.8 K (p < 0.05) and 3.8 K (p < 0.001) and maximum *VPD* differences of 0.2 kPa (p > 0.05) and 0.8 kPa (p < 0.05) in *F. sylvatica* and *P. menziesii*, respectively. Differences of *T_cuv_* (1.3°C and 0.7°C, respectively) and *VPD* (0.1 kPa and 0.1 kPa, respectively) were lower during early summer and during late summer, but due to low data availability no model could be fitted for this period (Figure 6 A-F). Strong difference of VPD and *T_cuv_* in midsummer resulted in 1.5-fold higher *g_s_* and *E* and twofold higher *A_net_* of sun leaves of *F. sylvatica* compared to shade leaves (p < 0.001), while no significant difference was found during early summer (Figure 6G-O). The deviation of sun and shade leaves of *F. sylvatica* during midsummer was mainly driven by an increase of *g_s_*, *A_net_* and *E* in the sun compared to early summer (p < 0.001), whereas leaf gas exchange in the shade remained similar. Contrarily, the difference of leaf gas exchange between sun and shade branchlets of *P. menziesii* remained constant during midsummer compared to early summer (p > 0.05), since no increase was observed for *g_s_, E* and *A_net_* despite higher *T_cuv_* and *VPD*. Stem water dynamics of both species resembled water fluxes of sun leaves with a 1.5-fold increase of maximum *J_s_* (p < 0.001) and no significant increase of *TWD* in *F. sylvatica* from early summer to midsummer, whereas only a non-significant increase of *J_s_* and a significant twofold increase of *TWD* (p < 0.001) of *P. menziesii* was observed (Figure 7).

**Figure 6:**
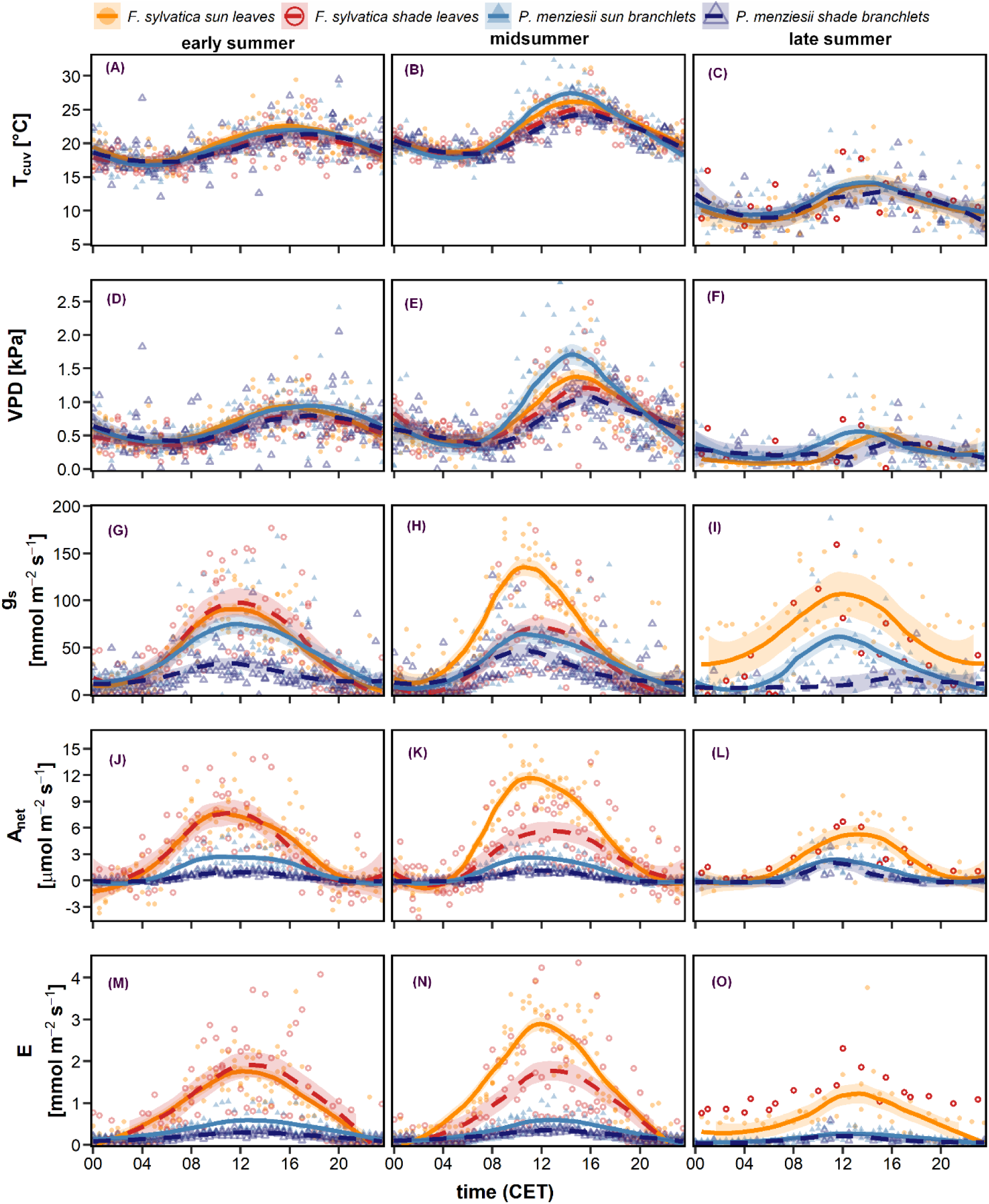
Mean diurnal courses of cuvette temperature (*T_cuv_*, A-C), cuvette vapor pressure deficit (*VPD*, D - F), stomatal conductance (*g_s_,* G-I), net carbon assimilation (*A_net_,* J-L) and water vapor transpiration (*E*, M-O) in the sun (solid lines, filled symbols) and shade canopy (dashed lines, empty symbols) of single leaves of *F. sylvatica* (n=3) and small branchlets of *P. menziesii* (n=3) during the three pre-defined periods. Points represent the hourly mean of the tree individuals during the respective period. Lines show a fitted generalized additive model, while shaded areas show the standard error of the model. Model fit of shade leaves of *F. sylvatica* in late summer is missing due to scarce data availability.

**Figure 7:**
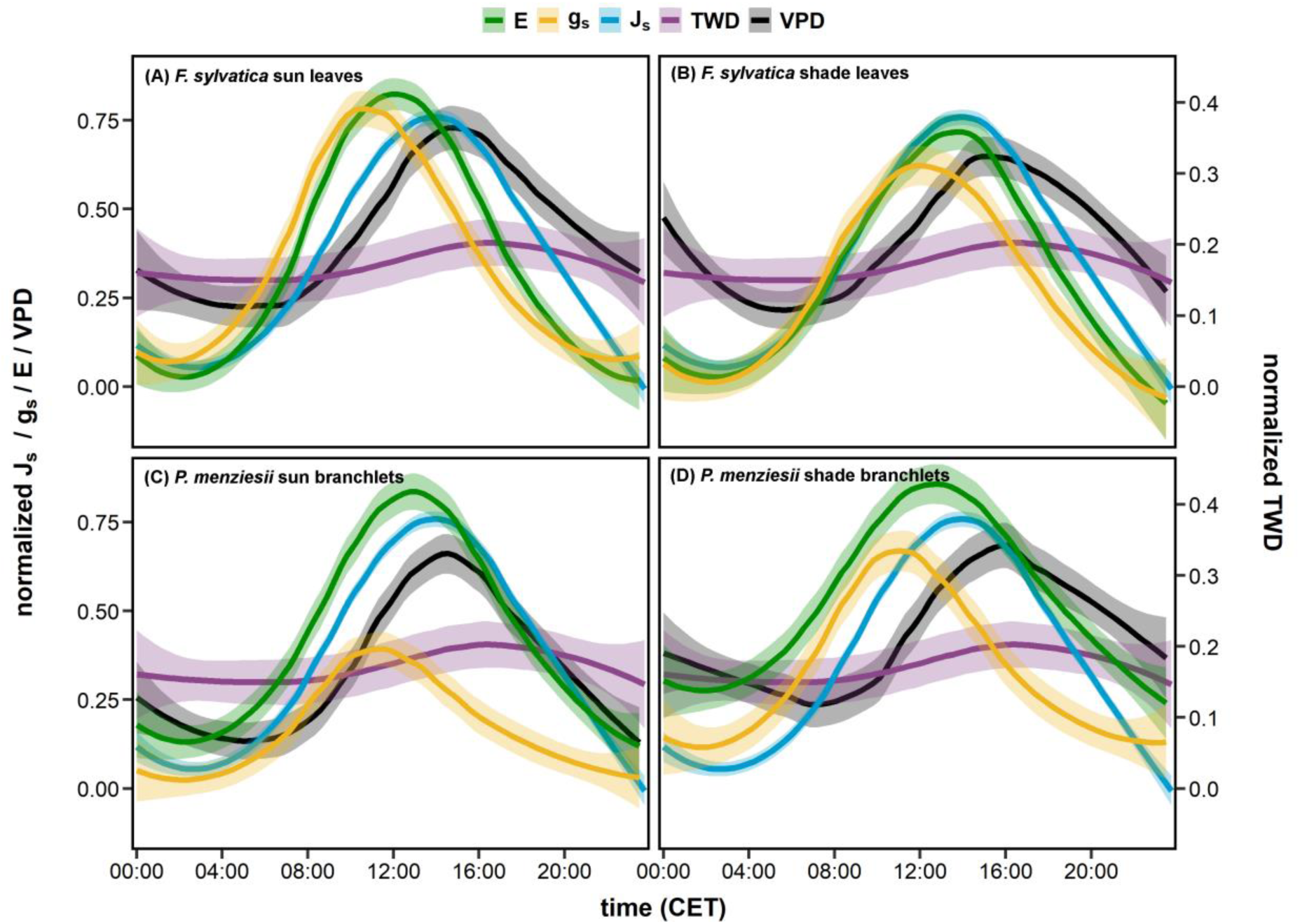
Diurnal courses of normalized sap flux density (*J_s_*), stomatal conductance (*g_s_*), transpiration (*E*) and tree water deficit (*TWD*) of sun (A) and shade (B) leaves of *F. sylvatica* (n=3) and of sun (C) and shade (D) branchlets of *P. menziesii* (n=3) during midsummer (see Table 1). Normalized values were calculated using the 99% quantiles of maximum values of the respective variable measured during the vegetation period of 2024. Shaded areas in the background show the standard error, while lines show the fitted curve of a generalized additive model.

A substantial time lag was detected between stem- and leaf-level water fluxes, particularly during midsummer (Figure 7): in sun leaves of *F. sylvatica*, maximum *A_net_* and *g_s_* were reached in the morning (*A_net_*: 10:00 am, *g_s_*: 10:30 am), followed one hour later by maximum *E* and 2.5 hours later by maximum *PPFD* and another 2 hours later by maximum *VPD*. In the shade foliage of *F. sylvatica*, maximum *g_s_* (11 am) was followed by a simultaneous peak of *A_net_* and *PPFD* (12.30 pm), 1.5 hours later by maximum *E* and another hour later by maximum *VPD*. *J_s_* and *TWD* peaked 1 and 1.5 hours after maximum *VPD* and 5 and 5.5 hours after maximum *g_s_* was reached in the canopy, respectively. In *P. menziesii*, maximum *g_s_* occurred first in in shade (10:30 am) and then in sun branchlets (11 am), followed by maximum *A_net_* (sun: 11:30 am, shade: 12 pm) and *PPFD* (12:30 am), and a simultaneous peak in *E* (1 pm). Maximum *VPD* was reached 3.5 (sun) to 5.5 hours (shade) after the peak of *g_s_* was recorded and another 2-3.5 hours later, already in the evening, maximum *J_s_* (6 pm) and *TWD* (6:30 pm) were observed. The same order was observed for both species during early summer, albeit time lags were slightly shorter (Supplementary Material Figure S4). Strong species differences were observed for the time lag until trees replenished their stem water reserves during midsummer: refilling in *F. sylvatica* was significantly faster (10:30 pm, 6 hours after maximum *J_s_*) than in *P. menziesii* (06:45 am, 13 hours after max *J_s_*).

## Discussion

In 2024, we established a sophisticated measurement system enabling continuous measurements of leaf gas exchange and campaign-based sampling of monoterpene emissions inside the tall tree canopy of a mixed temperate forest and combined it with stem-level water flux measurements. In *P. menziesii*, high VPD during midsummer was buffered in the shade and thus, only the sun canopy reduced stomatal conductance and monoterpene emissions in response to a period of gradual edaphic and atmospheric drying. In *F. sylvatica*, we found that shade foliage contributes significantly to whole-canopy photosynthetic carbon uptake and monoterpene emissions under overcast conditions, since net carbon assimilation, transpiration and emissions did not differ between the sun and shade canopy during overcast periods with low *T_air_*. We found substantial time lags between water fluxes in the stem and the canopy indicating a significant contribution of stored stem water reserves to leaf-level transpiration. In both species, our continuous measurement system revealed a high seasonal and day-to-day variability of VOC emissions, leaf gas exchange and stem-level water fluxes, which can provide a way forward in understanding coordination of whole-tree water and carbon fluxes.

### Microclimatic gradients shape within-canopy response to soil drying

In response to a dry-down period in midsummer, sun foliage of *P. menziesii* gradually decreased *g_s_* and increased *WUEi*, while the shade foliage was unaffected. Stomatal closure of *P. menziesii* resulted in a regulation of *J_s_*, *E* and *A_net_*, which did all not increase from early summer to midsummer despite more favorable weather conditions, a decline of *ψ_PD_*, *ψ_MD_* and *Δ_leaf_,* and an increase in *TWD*, which could not be fully replenished overnight (Figure 3, 4 & 7). Hence, the increasing water potential gradients in response to high atmospheric demand in the sun triggered stomatal closure to prevent excess loss of water and potentially protect stem water reserves and ensure sufficient cell turgor (Dumberger et al. 2025; Peters et al. 2025; Salomón and Camarero 2025). This is in line with findings from Schumann et al. (2024) and Paligi et al. (2025) who found a more isohydric response for *P. menziesii* than for *F. sylvatica* under drought, with early closure of stomata. Moreover, stomata of sun foliage generally respond faster to more water-limiting conditions in the soil, since they are exposed to higher levels of *PPFD*, *T_air_* and *VPD* (Niinemets et al. 2004b). Therefore, *g_s_* of sun foliage responds more sensitive to reductions in soil moisture, while shade foliage is more decoupled from soil water availability leading to dampened carbon uptake in the sun canopy under mild water stress (Wozniak et al. 2020). Microclimatic buffering of *T_air_* and *VPD* in the shade is assumed to reduce negative impacts of severe heat and drought stress on whole-canopy carbon uptake (He et al. 2018). Our findings show, that even under moderate soil drying, lower *VPD* in the shade of *P. menziesii* prevented shade foliage from the gradual, twofold reduction of *g_s_* observed in sun foliage and the subsequent regulation of *A_net_* and *E* despite warmer and sunnier conditions. However, absence of stomatal regulation also led to lower *WUEi* in shade foliage of *P. menziesii* and it is still uncertain how lower *WUE* of shade leaves will limit their contributions to canopy photosynthesis under severe long-term stress (Way and Pearcy 2012; Valladares et al. 2016). Lower *WUEi* in the shade is a result of microclimatic buffering and high residual *g_s_* under low *PPFD* ensuring high intercellular CO_2_ concentrations and thus, a quick activation of RuBisCo to efficiently exploit dynamic light environments (Way and Pearcy 2012; Campany et al. 2016). Future research with continuous leaf gas exchange data as presented in this study will be needed to better understand within-canopy dynamics of *g_s_* under severe stress conditions. However, from our data, we can show that under moderate soil and atmospheric drying lower *WUEi* of shade foliage has no negative impact on *A_net_* or *E*.

Simultaneous to decreasing *g_s_*, *ψ_leaf_* and *VWC* during midsummer, we also found decreased VOC emission rates of sun foliage in *P. menziesii*. Mild drought stress has been found to slightly increase monoterpene emissions, while severe drought stress drastically reduces emissions rates (Staudt et al. 2002; Wu et al. 2015; Haberstroh et al. 2018). We did not observe severe drought stress, but declining soil moisture did induce a reduction of *g_s_* and thereby a down-regulation of *E* and *J_s_*, thus we conclude that at least partially low emissions of *P. menziesii* during midsummer were elicited by decreasing soil moisture availability. *P. menziesii* comprises of specialized storage compartments, from which VOCs can be emitted under sufficient air temperature even if de novo synthesis rates are low (Lerdau et al. 1995; Joó et al. 2011). However, we also observed a clear seasonal decline in monoterpene emissions from July to September in *F. sylvatica*, which has also been observed by other studies (Pressley et al. 2004; Hakola et al. 2006; Holzke et al. 2006; Aalto et al. 2014). From our data, we cannot disentangle whether low monoterpene emissions of *P. menziesii* during midsummer resulted from decreasing *VWC*, decreased de novo synthesis, or from a seasonal decline. For future studies, the analysis of ẟ^13^C values of emitted compounds could improve our understanding of driving factors of seasonal VOC emissions (Ghirardo et al. 2010; Haberstroh et al. 2019; Daber et al. 2025).

We did not find a downregulation of *g_s_* in *F. sylvatica* during midsummer, but differences of *VPD* and *T_cuv_* between sun and shade leaves were also lower (Figure 6 B & E) and no strong reduction of *ψ_PD_* could be detected. Lower difference in *T_cuv_* and *VPD* between sun and shade foliage of *F. sylvatica* compared to *P. menziesii* were most likely related to the increase in *E* and *J_s_* and the subsequent transpirational cooling effect. *F. sylvatica* is known to respond with a more anisohydric, water-spending strategy to drying, maintaining transpiration even under slight water stress (Schumann et al. 2024; Paligi et al. 2025). However, during midsummer we observed that *g_s_* and *E* of sun leaves reached their peak values around 1.5 hours earlier than shade leaves, and three hours before highest *VPD* was recorded (Figure 7). This time lag was probably related to the higher radiation load in sun leaves and the subsequent faster increase of *VPD* during the morning hours. Thus, we conclude that, even though *g_s_* of more anisohydric *F. sylvatica* was not reduced during midsummer, *g_s_* still responded to rising *VPD* under high *PPFD*, yet not as drastically as observed in more isohydric *P. menziesii*.

### Shade leaves of F. sylvatica benefit from overcast skies

During early and late summer, we found similar *g_s_*, *A_net_* and *E* in the sun and shade leaves of *F. sylvatica* under overcast skies with frequent precipitation events, whereas sun leaves clearly exceeded shade leaves in *g_s_*, *A_net_* and *E* during midsummer (Figure 3 & 4). Higher maximum photosynthetic and electron transport capacity of sun leaves optimize them to efficiently utilize direct sunlight, while shade leaves are better adapted to exploit diffuse light environments (Niinemets 2023). Increasing the fraction of diffuse light to direct light in modelling simulations resulted in 10-50% higher canopy assimilation rates, which were mainly related to higher contributions of the shade foliage (Meir et al. 2002; Dai et al. 2004). Diffuse light can better penetrate the canopy and increase the radiation load on otherwise light-limited shade leaves and has been found to contribute around 64% to global gross primary productivity despite contributing only 54% to total available light (Knohl and Baldocchi 2008; Zhou et al. 2021). Global modelling showed that on average shade canopy contributes to ∼48% of gross primary productivity in broadleaved deciduous forests (He et al. 2018) and that big-leaf models underestimate canopy productivity between 20% and 70% by neglecting differing behavior of shade leaves (Meir et al. 2002; Dai et al. 2004; Sprintsin et al. 2012). In *F. sylvatica*, only around 20% of total leaf area index is comprised of sunlit leaves (Scartazza et al. 2016), which would imply that in our study around 80% of whole-canopy assimilation was achieved by shade foliage during early summer and late summer.

Furthermore, we found that monoterpene emissions of the sun leaves of *F. sylvatica* were usually higher than those of the shade leaves on sunny days, but that this difference diminished on cloudy days due to higher emissions of the shade leaves than under sunny conditions (Figure 5). Using an eddy covariance system in a mixed forest ecosystem, Laffineur et al. (2013) also found higher VOC emissions under cloudy than under clear-sky conditions, which was probably related to the higher contribution of shade leaves to emissions on these days. Since *F. sylvatica* does not comprise of specialized VOC storage structures, emissions are mainly dependent on *PPFD*, *T_air_* and substrate availability from photosynthesis (Niinemets et al. 2004a; Dindorf et al. 2006; Van Meeningen et al. 2016; Bourtsoukidis et al. 2024). Therefore, we assume that better penetration of diffuse light into the canopy and resulting higher photosynthetic activity fostered monoterpene emissions of the shade leaves.

We did not find the same pattern in the shade foliage of *P. menziesii*, where *g_s_*, *A_net_* and *E* were lower than in the sun foliage throughout the entire study period (Figure 3). Most likely, different canopy architecture of our coniferous and deciduous species led to the observed differences, since foliage clumping is higher in *P. menziesii* than in *F. sylvatica* which distributes light more evenly within the canopy (Stenberg 1998). Such penumbral effects need to be considered regarding our gas exchange measurements, since whole branchlets of *P. menziesii* were enclosed into the cuvettes and calculations were done with projected leaf area. For better comparability, development of a needle cuvette system as deployed for *F. sylvatica* would be useful (Frey et al. 2025a), but remains challenging due to the needle arrangement around the branchlets and small leaf area of a single needle compared to a single leaf of *F. sylvatica*.

### Seasonal and spatial variability of monoterpene fluxes

Both VOC emissions and leaf gas exchange were in the range of previous studies investigating our two species in campaign-wise measurements (Lerdau et al. 1995; Pressley et al. 2004; Dindorf et al. 2006; Holzke et al. 2006; Joó et al. 2011; Šimpraga et al. 2011; Laffineur et al. 2013; Scartazza et al. 2016; Durand et al. 2022), but clearly lack diurnal and day-to-day variability and/or seasonal effects, which we were able to detect with our continuous measurement system. Our data can crucially enhance parametrization of current upscaling approaches and help to improve model uncertainties elicited by microclimatic variability (Wozniak et al. 2020), foliage clumping effects (Chen et al. 2012), seasonal dynamics (Keenan et al. 2009; Guenther et al. 2012; Grote et al. 2013; Chang et al. 2018), and plant hydraulic constraints (Peltoniemi et al. 2012; Niinemets 2023).

### Stomatal aperture is tightly coordinated between stem-level water supply and leaf-level water demand

We observed substantial time lags between diurnal stem- and leaf-level water fluxes, particularly during midsummer (Figure 7). In *P. menziesii*, refilling of stem water reserves lasted the entire night which was most likely related to reduced soil water supply and lower hydraulic conductance of coniferous xylem tracheids compared to ring-porous vessels of *F. sylvatica* (Berdanier et al. 2016). Differences in hydraulic conductance between both species were also visible in the time lag between maximum canopy *VPD* and maximum *J_s_* in the stem, which was only one hour in *F. sylvatica* but three hours in *P. menziesii*. Moreover, since in both species maximum *E* occurred earlier than maximum *J_s_* and simultaneously we observed an increase in *TWD*, water demand during peak *E* was partially supplied by stored stem water. This is in line with previous studies which observed that stored stem water contributed 9-16% in *F. sylvatica* (Köcher et al. 2013; Leuschner et al. 2024) and 20-25% in *P. menziesii* (Phillips et al. 2003; Cermak et al. 2007) to total water consumption. This also indicates that during the afternoon, when peak *J_s_* occurred, transported water was partially used to refill depleted stem water reserves along the stem, since demand by the crown was already reduced during this time of the day. Water withdrawal from the stem during morning hours to meet transpirational demand by the crown and refilling of depleted tissue by *J_s_* during the afternoon was also observed by (Köcher et al. 2013) for five broadleaved temperate species.

Generally, in both species, we observed a temporal relationship between peak values of *J_s_* and *VPD*, *E* and *PPFD*, and *g_s_* and *A_net_.* Interestingly, in both species, maximum *g_s_* peaked close to maximum *A_net_* and occurred four to six hours before maximum *VPD* was reached indicating a tight stomatal control in response to *VPD*. We see three potential explanations for stomatal regulation: (1) stomatal aperture is optimized for maximum carbon gain and maximum *A_net_* happened already in the morning (Henry et al. 2019; Deans et al. 2020; Franklin et al. 2023), (2) reduced *VWC*, delayed stem water refilling and resulting decline in *ψ_PD_* led to early stomatal closure (Peters et al. 2025), (3) or a species-specific threshold of *VPD* (VPD > 0.5 kPa) elicits stomatal regulation independent of maximum *VPD* (Oren et al. 1999; Novick et al. 2016; Grossiord et al. 2020), as it has also been observed for the onset of radial growth (Zweifel et al. 2021). However, we showed that maximum *g_s_* occurred earlier during midsummer than during early summer and also earlier in the sun than in the shade foliage of both species. Therefore, we conclude that stomatal regulation represents a fine-tuned coordination between *ψ_leaf_*, *VPD* and *A_net_* at the leaf level and *TWD_min_* and *VWC* at the stem base and cannot be explained by one factor alone.

## Conclusion

Continuous measurements of leaf gas exchange in the sun and shade canopy of *F. sylvatica* and *P. menziesii* revealed strong seasonal dynamics of stomatal regulation as well as of VOC emissions. More isohydric *P. menziesii* responded sensitive to increasing *VPD* with a reduction of stomatal conductance and VOC emissions in the sun branchlets and a downregulation of sap flux density, while shade branchlets did not respond due to microclimatic buffering of *VPD*. Contrarily, sun leaves of more anisohydric *F. sylvatica* profited from higher air temperature and *VPD* in midsummer, while shade leaves were limited by low light availability. Under overcast skies, shade leaves of *F. sylvatica* benefitted from better penetration of diffuse light into the canopy and therefore showed similar carbon assimilation and VOC emissions to sun leaves. Time lags between stem- and leaf-level water fluxes were most pronounced during midsummer, when air temperature and *VPD* were highest, and up to 20 hours were needed to replenish stem water reserves. Pronounced seasonal courses of stomatal regulation processes and VOC emission patterns, which varied between the sun and shade canopy, demonstrated that continuous leaf gas exchange data in different canopy layers combined with stem-level flux measurements are indispensable for a better understanding of whole ecosystem water and carbon fluxes under stressed and non-stressed conditions.

## Acknowledgements

We would like to thank Lennart Nettler and Phyllis Lua-Mellmann for assistance with sensor installations and maintenance and L. Erik Daber and Tim Ehrhardt for the support during the construction and installation of the measurement system. We thank Mirjam Meischner for the fruitful discussions of the on-site development of the measurement system and on calculations of leaf gas exchange data. We also gratefully acknowledge permission from the city of Ettenheim to set-up our field site in their city forest.

## Author Contributions

Leading author: Stefanie Dumberger: experimental setup, data collection, analysis, and interpretation, manuscript writing

Yasmina Frey: sensor development, data collection, manuscript revision

Clara Stock: experimental setup, data collection and interpretation, manuscript revisio

Sophie Wehlings-Schmitz: data collection and analysis of VOC samples

Delon Wagner: electrical setup of the measurement equipment, software programming

Kathrin Kühnhammer: electrical setup and maintenance of the measurement equipment

Lea Dedden and Markus Weiler: data acquisition of soil moisture sensors, manuscript revision

Markus Sulzer and Andreas Christen: data acquisition of meteorological data, manuscript revision

Jürgen Kreuzwieser: analysis and interpretation of VOC emissions, manuscript revision

Ulrike Wallrabe: study conception, manuscript revision, supervision

Christiane Werner: study conception, data interpretation, manuscript revision, supervision

Simon Haberstroh: study conception, data analysis and interpretation, manuscript revision

## Open Research

The data that support the findings of this study are available from the corresponding author upon reasonable request.

