## Supplementary Material for "Seasonal dynamics and sun/shade heterogeneity of leaf gas exchange and VOC emissions inside a tall temperate forest canopy"

### Calculation of leaf gas exchange

The following formulas were used to calculate transpiration (E, Formula S1), carbon assimilation ($A_{net}$, Formula S2), stomatal conductance for water vapour ($g_s$, Formula S3) and plant discrimination for $^{13}C$ ($\Delta_{plant}$, Formula S4) of the enclosed leaves and branchlets (von Caemmerer and Farquhar 1981; Evans et al. 1986):

$$E = \frac{u}{s} \times \frac{(w_o - w_e)}{(1 - w_o)} \qquad \text{(Formula S1)}$$

where E is the transpiration rate in mmol $m^{-2}$ $s^{-1}$, u is the molar flux in mol $s^{-1}$ of air through the cuvette, s is the leaf area in $m^2$, $w_o$ is the mass fraction of water vapour exiting the cuvette [mol $mol^{-1}$] and $w_e$ is the mass fraction of water vapour entering the cuvette [mol $mol^{-1}$].

$$A_{net} = \frac{u}{s} \times \left(\frac{1 - w_e}{1 - w_o}\right) \times (c_e - c_o) - E \times c_e \qquad \text{(Formula S2)}$$

where $A_{net}$ is the $CO_2$ assimilation rate [µmol $m^{-2}$ $s^{-1]}$, $c_o$ is the mass fraction of $CO_2$ exiting the cuvette [mol $mol^{-1}$] and $c_e$ is the mass fraction of $CO_2$ entering the cuvette [mol $mol^{-1}$].

$$g_s = \frac{E \times \left(1 - \frac{w_i + w_a}{2}\right)}{(w_i - w_a)} \qquad \text{(Formula S3)}$$

where $g_s$ is stomatal conductance for water vapour in mmol $m^{-2}$ $s^{-1}$, $w_i$ is the mass fraction of water vapour inside the leaf [mol $mol^{-1}$] and $w_a$ is the mass fraction of water vapour outside the leaf [mol $mol^{-1}$].

$$\Delta_{leaf} = \frac{1000 \times (\frac{c_e}{(c_e - c_o)} \times (\delta_o - \delta_e)}{(1000 + \delta_o - (\frac{c_e}{(c_e - c_o)} \times (\delta_o - \delta_e)))} \qquad \text{(Formula S4)}$$

where $\Delta_{leaf}$ is the apparent leaf $^{13}C$ discrimination during photosynthesis [‰], $c_e$ is the mass fraction of $CO_2$ entering the cuvette [mol $mol^{-1}$], $c_o$ is the mass fraction of $CO_2$ exiting the cuvette [mol $mol^{-1}$], $\delta_e$ is the $\delta^{13}C$ signature [‰] of air entering the cuvette and $\delta_o$ is the $\delta^{13}C$ signature [‰] of air exiting the cuvette.

Thereafter, calculated values of $A_{net}$, E and $g_s$ were used for calculations of water use efficiency (WUE, Formula S5) and intrinsic water use efficiency (WUEi, Formula S6):

$$\mathrm{WUE} = \frac{\mathrm{A_{net}}}{\mathrm{E}} \qquad \text{(Formula S5)}$$

where WUE is water use efficiency in µmol C $mmol^{-1}$ $H_2O^{-1}$.

$$\mathrm{WUEi} = \frac{\mathrm{A_{net}}}{\mathrm{g_s}} \qquad \text{(Formula S6)}$$

where WUEi is the intrinsic water use efficiency in µmol C $mmol^{-1}$ $H_2O^{-1}$.

BVOC fluxes were calculated using Formula S7:

$$BVOC\ flux\ = \frac{\frac{(c_o - c_e) \times \frac{u_e}{u_o}}{s}}{M} \qquad (Formula\ S7)$$

where BVOC flux is the emission flux of BVOCs [nmol $m^{-2}$ $h^{-1}$], $c_o$ is the concentration of BVOCs exiting the cuvette [ng], $c_e$ is the concentration of BVOCs entering the cuvette [ng], $u_e$ is the air flux entering the cuvette [ml $min^{-1}$], $u_o$ is the air flux drawn from the cuvette [ml $min^{-1}$], s is the leaf area [$m^2$] inside the cuvette and M is the molar Mass [g $mol^{-1}$] of the specific compound.

### **Additional Figures**

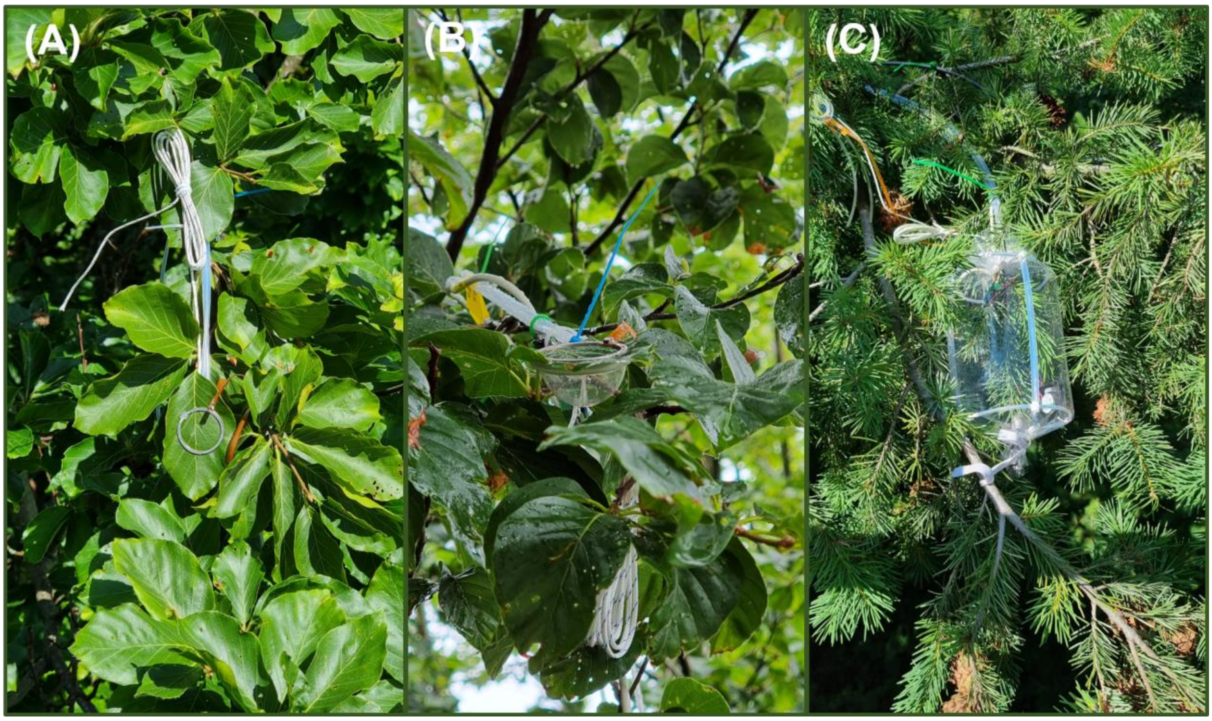


Figure S1: Images of the employed leaf enclosures for gas exchange measurements in the forest canopy. Shown are a top view (A) and a side view (B) of the novel ECOvettes (Frey et al. 2025) used for single leaves of *F. sylvatica* as well as a custom-built cuvette made of FEP film and PFA tubing (C) used for small branchlets of *P. menziesii*.

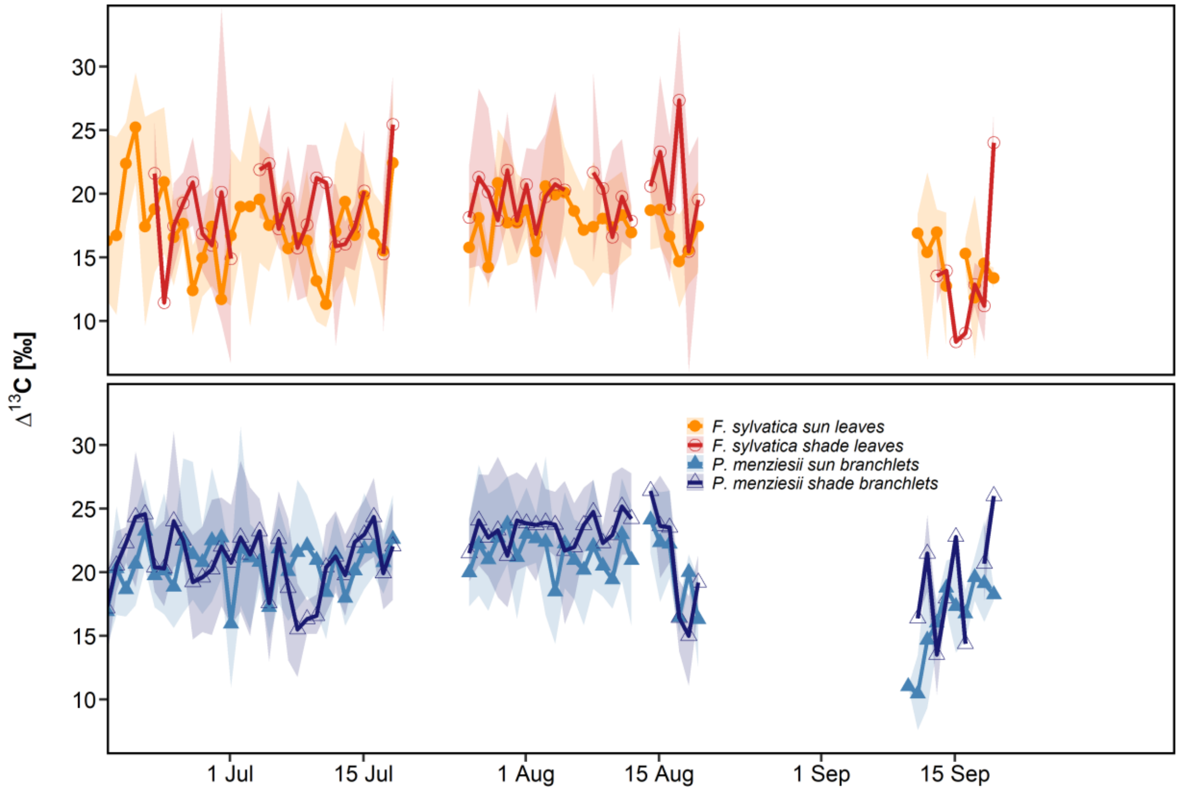


Figure S2: Daily mean leaf $^{13}C$ discrimination ($\Delta^{13}C$) of sun-exposed (bright color) and shade exposed (dark color) leaves of *F. sylvatica* (upper panel, n=3) and branchlets of *P. menziesii* (lower panel, n=3). Shaded area in the background indicates the daily maximum and daily minimum. $\Delta^{13}C$ averaged over all tree individuals.

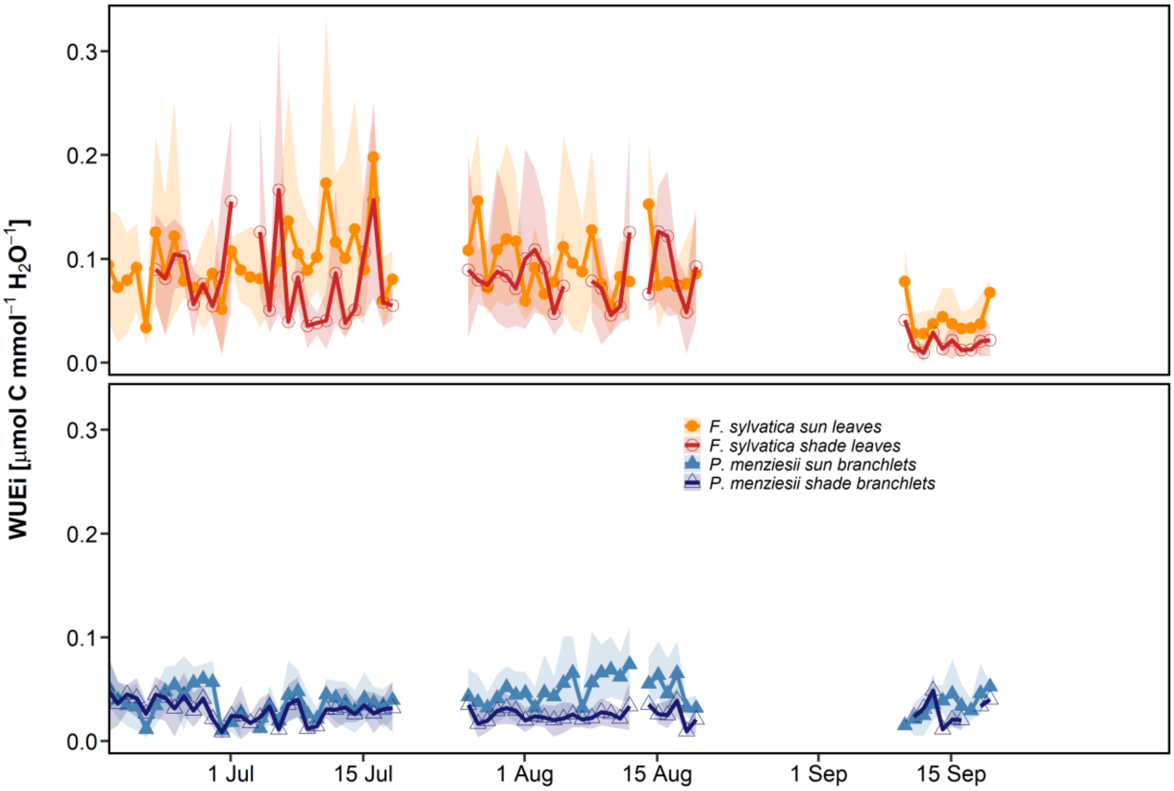


Figure S3: Daily mean intrinsic water use efficiency (*WUEi)* of sun-exposed (bright color) and shade exposed (dark color) leaves of *F. sylvatica* (upper panel, n=3) and branchlets of *P. menziesii* (lower panel, n=3). Shaded area in the background indicates the daily maximum and daily minimum *WUEi* of all tree individuals.

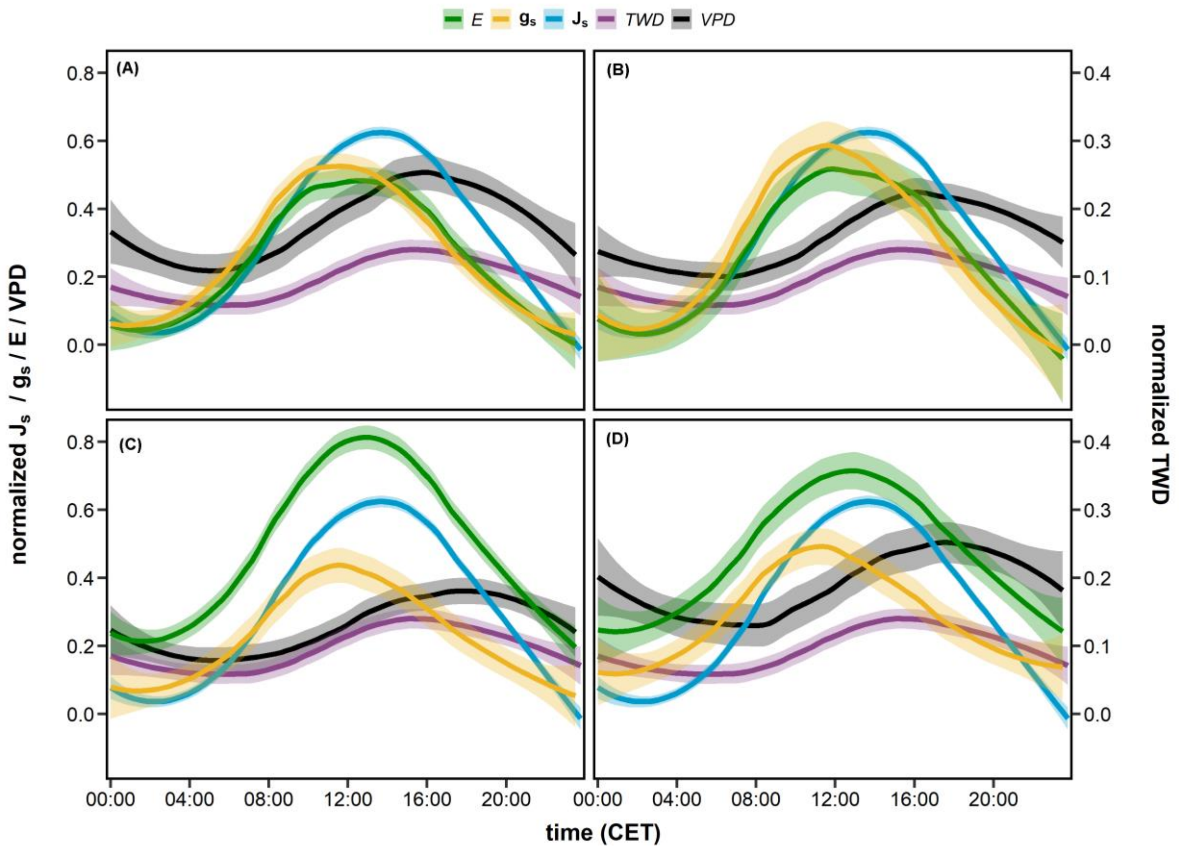


Figure S3: Diurnal courses of normalized sap flux density ($J_s$), stomatal conductance ($g_s$), transpiration (*E*) and tree water deficit (*TWD*) of sun (A) and shade (B) leaves of *F. sylvatica* (n=3) and of sun (C) and shade (D) branchlets of *P. menziesii* (n=3) during early summer (see Table 1). Normalized values were calculated using the 99% quantiles of maximum values of the respective variable measured during the vegetation period of 2024. Shaded areas in the background show the standard error, while lines show the fitted curve of a generalized additive model.
